# A Foundation Model for the Cancer Genome

**DOI:** 10.64898/2026.05.27.728319

**Authors:** John-William Sidhom, Alex S. Baras, Olivier Elemento, Manish A. Shah

**Affiliations:** Division of Hematology & Medical Oncology, Weill Cornell Medical College, New York, NY, USA; Department of Pathology, The Johns Hopkins University, Baltimore, MD, USA; Englander Institute for Precision Medicine, Weill Cornell Medicine, New York, NY, USA; Department of Systems and Computational Biomedicine, Weill Cornell Medicine, New York, NY, USA; Institute for Computational Biomedicine, Weill Cornell Medicine, New York, NY, USA; Center for Advanced Digestive Care, NewYork-Presbyterian, New York, NY, USA

## Abstract

Cancer is a disease of the genome, in which somatic mutations and copy-number alterations determine tumour identity, clinical behaviour, and response to therapy. Consortium-scale sequencing has profiled hundreds of thousands of tumours,^1,2^ yet clinical interpretation still proceeds one alteration at a time against hand-curated knowledgebases,^3,4^ often ignoring co-occurring alterations and the genome-wide copy-number pattern. Self-supervised foundation models pretrained on unlabelled corpora^5^ have produced transferable representations in adjacent biological domains^6–8^ by learning joint structure across many features, yet no comparable model exists for the cancer genome. Here we present TESSERA (Tumour Embeddings via Self-Supervised Encoding and Reconstruction of Alterations), a foundation model for the cancer genome; we pretrain it on somatic single-nucleotide variants and copy-number segments through masked-token reconstruction within each modality and a contrastive objective across modalities. A single representation, produced once and reused without retraining, supports variant pathogenicity prediction, pan-cancer tumour typing, unsupervised molecular subtyping, prognostic stratification, and counterfactual treatment-effect estimation that yields predictive chemotherapy-selection biomarkers in real-world cohorts. These biomarkers are interpretable: each surfaces the co-occurring alterations underlying the prediction, exposing biology that single-gene rules miss. In metastatic colorectal cancer, where the FOLFOX-vs-FOLFIRI choice is currently guided by toxicity rather than tumour biology, the model uncovers a candidate predictive biomarker: a three-feature rule (TP53^+^/KRAS^+^/17p^−^) selecting patients who derive substantially greater benefit from FOLFOX than FOLFIRI.

## Introduction

Somatic genotype is increasingly central to how oncologists diagnose, prognose, and treat cancer. BRAF V600E melanomas are matched to dabrafenib, EGFR-mutant lung adenocarcinomas to osimertinib, BRCA-deficient breast and ovarian tumours to PARP inhibitors, HER2-amplified breast and gastric tumours to trastuzumab, and an expanding catalogue of single-gene biomarkers steers clinical decision-making across several solid tumours. Consortium-scale sequencing was launched in the hope of discovering more such markers and uncovering druggable pathways: the TCGA Pan-Cancer Atlas^1,9^ characterised somatic alterations across more than thirty tumour types,^10,11^ the PCAWG whole-genome collection extended this to non-coding variation,^12^ and the AACR Project GENIE consortium has aggregated panel-based clinical NGS data from multiple cancer centres.^13^ Despite this scale of data, predictive biomarkers in routine oncology practice remain almost exclusively single-gene, and clinical sequencing reports interpret each alteration separately, even as the same tumour carries dozens to hundreds of co-occurring alterations.

The prevailing paradigm for interpreting genomic alteration data is gene-by-gene. Individual somatic variants are annotated against hand-curated knowledgebases such as ClinVar for germline pathogenicity^3^ and OncoKB for oncogenicity and actionability,^4^ and classified using the AMP/ASCO/CAP consensus tiers.^14^ Driver-gene discovery relies on statistical scans over consensus mutation calls.^15^ These frameworks encode substantial domain expertise but are locally evaluative: each variant is interpreted in isolation against a curated reference, without using the context of co-occurring alterations, sample-level mutational signatures, or the genome-wide pattern of copy-number change. The mismatch between the scale of available data and the granularity of interpretation motivates approaches that learn a joint representation of a tumour’s genome directly from the data.

Self-supervised foundation models now produce transferable representations in adjacent biological domains. In language and vision, models pretrained through masked-token^16^ or contrastive^17^ objectives produce representations that transfer to many tasks without per-task retraining.^5^ The same approach is now used in models of protein sequence and structure (ESM-2, AlphaFold),^6,18^ germline DNA (Enformer),^19^ single-cell transcriptomics (Geneformer),^7^ and cancer histopathology (UNI).^8^ In each case, a single pretrained model has replaced task-specific classifiers and supports many downstream applications.

These successes do not directly transfer to the cancer genome. Models such as AlphaMissense,^20^ Enformer,^19^ and AlphaGenome^21^ are trained on reference and germline data, not on the somatic alterations cancer selects, and none represent copy-number alterations, a defining feature of cancer mutagenesis. Applied to cancer somatic mutations, AlphaMissense and related next-generation pathogenicity predictors perform commendably on general driver detec-tion but underperform cancer-specific methods and cannot resolve cancer-type-specific drivers;^22^ they also fail to identify gain-of-function variants,^23^ a major class of cancer-selecting alterations. A foundation model for the cancer genome therefore needs to be pretrained on the cancer genome itself, not adapted from reference- or germline-trained predecessors.

No analogous model exists for the cancer genome. Unlike research-grade assays such as single-cell transcriptomics, somatic sequencing is now routine at the time of cancer diagnosis, since many therapies require a specific genomic indication for use; the clinical data substrate is therefore growing at the pace of standard oncology care. The methods currently applied to this data remain narrow and single-task. Pan-cancer tumour-type classifiers^24,25^ are an academically tractable benchmark that does not by itself address the prognostic and predictive questions driving clinical decisions. Pan-cancer multi-modal survival models,^26^ unsupervised mutational-signature decompositions over single-nucleotide variation,^10^ and biomarker-specific chemotherapy-outcome predictors each target a single objective. None produces a pretrained, transferable representation of a tumour’s joint somatic-variant and copy-number-alteration profile that can be applied, without task-specific retraining, to the range of downstream problems in clinical oncology.

Here we introduce TESSERA (Tumour Embeddings via Self-Supervised Encoding and Reconstruction of Alterations), a foundation model for the cancer genome. TESSERA is jointly pretrained on somatic single-nucleotide variants and copy-number segments from the TCGA Pan-Cancer Atlas through masked-token reconstruction within each modality and a cross-modal contrastive objective, producing a sample-level representation that aligns variant and copy-number views of the same tumour. This places the cancer genome alongside protein sequence (ESM-2),^18^ single-cell transcriptomics (Geneformer),^7^ and tissue histology (UNI)^8^ as a domain served by a single self-supervised representation. The same representation, produced once and reused without retraining, supports oncology tasks at three biological scales: at the variant scale, clinical pathogenicity prediction; at the sample scale, tumour-type classification and unsupervised recovery of established molecular subtypes; and at the patient scale, prognostic risk stratification and counterfactual treatment-effect estimation. The learned features transfer from whole-exome pretraining to targeted-panel clinical-sequencing cohorts without fine-tuning, and, within the prognostic-versus-predictive framework for clinical biomarkers,^27^ produce a genomic treatment-selection score that satisfies the direction-reversal criterion of a predictive biomarker in real-world first-line metastatic chemotherapy.

## Results

### Masked-token SNV foundation model

We first pretrain a self-supervised encoder over somatic single-nucleotide variants (SNVs) on variants from ∼10,000 TCGA cancer samples (Fig. 1a). Following the masked-language-modelling approach, the model predicts masked reference and alternate alleles given the variant’s genomic position and two complementary forms of context: a local attention module over flanking genomic sequence (1bp, 10bp or 25bp on each side) and a global attention module over other variants within the same sample. We compared seven architectural variants with progressively richer context: a baseline with no context; local-only attention at each of the three flanking widths; and global (local + global) at the same three widths.

**Figure 1.**
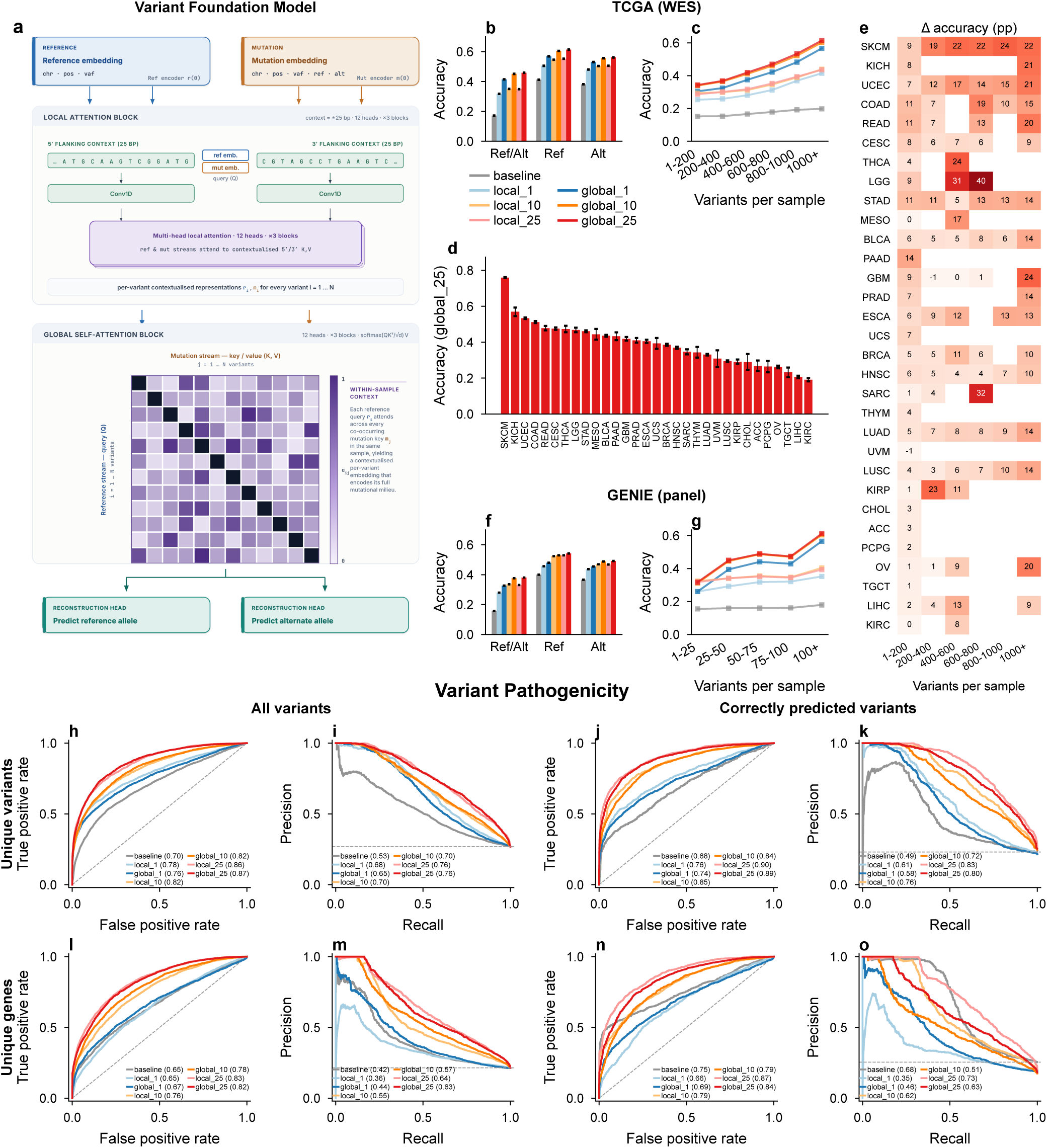
Somatic-variant foundation model for masked-token prediction and pathogenicity. **a,** Variant foundation model architecture. A reference embedding (blue; chromosome, position, variant allele frequency) and a mutation embedding (orange; same features plus reference/alternate alleles) are maintained as parallel streams. A local attention module cross-attends each stream to 5’ and 3’ flanking context processed through 1D convolutional layers. A global attention module uses the reference stream as query (Q) and the mutation stream as key/value (K, V), letting each variant position attend to mutations at other positions while masked from its own alleles. The attended query predicts the masked reference and alternate alleles. **b–e,** Per-variant masked-token prediction on TCGA WES. **b,** accuracy across architectures stratified by prediction type (Ref/Alt both correct; Ref only; Alt only), comparing baseline (no context), local attention with 1/10/25bp flanking sequence, and global attention at the same widths; error bars are 95% bootstrap CIs (n = 1,000). **c,** accuracy stratified by per-sample variant burden. **d,** accuracy by cancer type across 31 TCGA solid tumours, for the global-attention 25bp model (global_25). **e,** heatmap of the absolute accuracy improvement (percentage points) from global (25bp) over local-only (25bp) attention, stratified by cancer type (rows) and variant-burden bin (columns). **f–g,** Cross-platform transfer to the independent GENIE panel-sequencing cohort: panel (f) is the per-variant accuracy comparison of panel (b), panel (g) is the burden stratification of panel (c). **h–o,** Variant pathogenicity prediction on ClinVar, reported as ROC (panels h, j, l, n) and precision-recall (panels i, k, m, o) curves across four evaluation strategies: unique-variant split on all variants (h–i) and restricted to variants whose masked reference and alternate alleles were correctly reconstructed during pretraining (j–k); gene-level split on all variants (l–m) and restricted to correctly-reconstructed variants (n–o). Models compared match panel (b). All curves are means across evaluation folds. AUC and AP values indicated in legends.

Masked-token accuracy improved with genomic context (Fig. 1b). Global attention achieved 45.9% accuracy at jointly predicting both reference and alternate alleles (Ref/Alt) with ±25bp context, compared with 17.2% for a context-free baseline; most of the gain was captured by local sequence context alone, with global attention (which adds variant-variant interactions) providing a smaller but consistent additional improvement. Accuracy was higher for reference-only or alternate-only prediction than for joint Ref/Alt prediction, reflecting the harder task of simultaneously learning the reference genome and variant-specific mutational patterns.

Masked-token accuracy rose with sample variant burden (Fig. 1c), with substantially larger gains for global-attention than local-only models, confirming that within-sample variant patterns inform prediction. At the sample level, higher burden produced progressively tighter accuracy distributions and higher median accuracy (Supplementary Figure 1). Across cancer types (Fig. 1d) the highest accuracy was observed in SKCM, consistent with its strong UV mutational signature. A per-cohort heatmap (Fig. 1e) plots the absolute accuracy improvement from global over local attention; it separates intrinsic signature strength from variant-count effects: global attention provides substantially larger improvements in tumours with strong mutational processes (notably SKCM and UCEC), and these improvements grow with variant burden. Applied to an independent GENIE panel-sequencing cohort spanning multiple cancer centres, the TCGA-trained models retained the same ordering and scale of performance (Fig. 1f–g), indicating that the learned variant representations capture generalisable genomic properties that transfer across sequencing technologies, institutions, and patient populations.

### Variant pathogenicity prediction

We next evaluated TESSERA’s SNV submodel for variant pathogenicity prediction using ClinVar annotations. Flanking sequence composition influences mutational signatures through trinucleotide context, codon position and resulting amino acid changes, and proximity to splice sites, so a representation that encodes local sequence context should also predict pathogenicity.

We trained and evaluated multiple model variants with different sequence context lengths, measuring both ROC AUC and average precision (AP). On unique variants unseen during training (Fig. 1h–i), local sequence context drove most of the performance improvement. Baseline models with no context achieved ROC AUC of 0.705, while models with local attention incorporating 25bp flanking sequence reached 0.865. Global-attention models, which additionally learn variant-variant interactions, reached 0.869, a small additional gain over local context alone. As expected, the pathogenicity of a variant is largely a variant-level property determined by local sequence context, not by other variants in the sample. Precision-recall curves (Fig. 1i) followed the same pattern.

We then asked whether the model’s masked-token reconstruction accuracy correlates with pathogenicity prediction performance, restricting the evaluation to variants whose masked reference and alternate alleles the model had correctly reconstructed during pretraining (Fig. 1j–k). Performance improved substantially on this filtered set: variants the model can reconstruct at the sequence level are also those where its pathogenicity predictions are most reliable, so masked-token reconstruction accuracy works as a per-variant confidence filter on the pathogenicity call.

To test generalisation more strictly, we then held out test variants from genes completely absent during training (Fig. 1l–m). This gene-level deduplication tests whether the model learns transferable sequence patterns rather than memorising gene-specific associations. Performance decreased slightly on this harder task, with baseline models reaching 0.653 AUC (vs. 0.705 on unique variants) and local attention models with 25bp context reaching 0.828 (vs. 0.865), but remained strong, indicating that the learned sequence-level patterns capture properties of variant pathogenicity independent of gene identity. Restricting to variants the model correctly reconstructed at pretraining (Fig. 1n–o) again sharpened performance, so masked-token reconstruction accuracy works as a confidence filter even when generalising to novel genes. Performance metrics across all evaluation strategies are provided in Supplementary Figure 2.

### Copy-number foundation model

We then trained a CNA foundation model on TCGA ABSOLUTE copy-number segments under a dual-task objective: continuous segment-mean regression and binary loss-of-heterozygosity (LOH) classification. The model (Fig. 2a) separates positional features from segment-mean and LOH values to prevent leakage, and allows each segment to attend to every other segment in the sample via self-attention with diagonal masking. We compared configurations with 0, 1 and 2 self-attention blocks.

**Figure 2.**
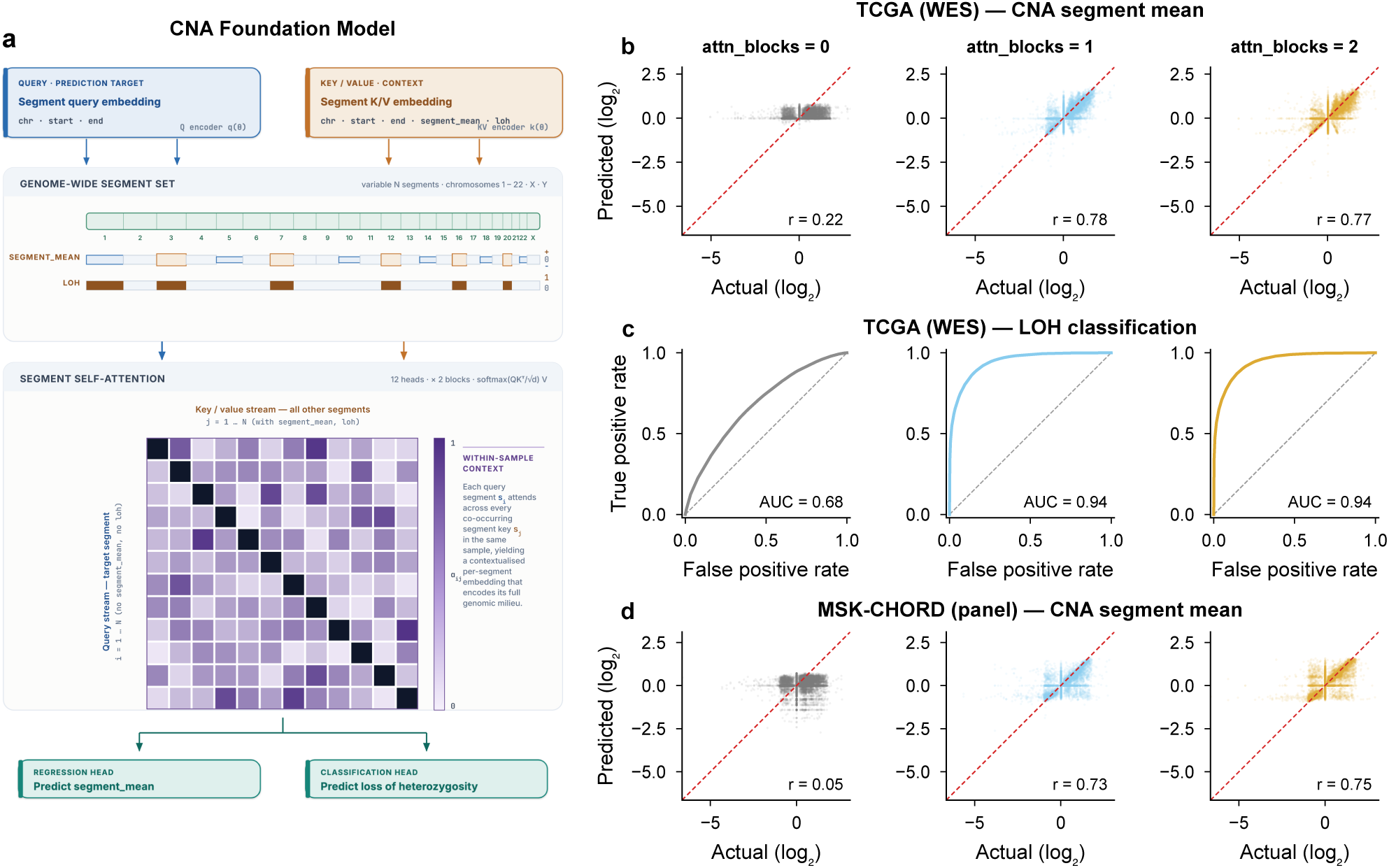
Copy number alteration prediction through self-attention on CNA segments. **a,** CNA foundation model architecture. The model maintains two embedding streams: a query embedding (blue) containing chromosome, start, end, and segment length, and a key/value embedding (orange) additionally containing segment mean and LOH status. In the self-attention module, the query embedding excludes segment mean and LOH to prevent data leakage, each segment cannot see its own values when making predictions. Segments attend to other segments’ copy-number and LOH states through self-attention with diagonal masking (preventing self-attention). The attended query is used to predict segment mean (regression) and LOH status (classification). **b,** Scatter plots of predicted versus actual segment mean (log2 copy-number ratio) on held-out TCGA validation segments. Models compared: baseline with no self-attention (0 attention blocks), and models with one and two self-attention blocks. Red dashed line indicates perfect prediction (y=x). Pearson correlation values shown in panel titles. **c,** ROC curves for binary LOH classification across attention block configurations. Curves show true positive rate versus false positive rate. AUC values indicated for each model: baseline (0 attention blocks), and models with one and two attention blocks. **d,** Scatter plots of predicted versus actual segment mean on independent MSK-CHORD panel-sequenced data. CNA segments were filtered to panel gene regions (MSK-IMPACT panels) and MSK-CHORD segment means were quantile-normalised onto the TCGA training segment mean distribution (rank-mapping) prior to inference, re-expressing MSK values at the TCGA amplitude scale while preserving each sample’s relative segment ordering. Models trained on TCGA WES data without LOH prediction (NoLOH variant), evaluated on MSK-CHORD. Same attention block configurations as panel (b). Pearson correlation values shown in panel titles.

Segment-mean prediction improved sharply with inter-segment attention (Fig. 2b): the baseline (0 blocks) achieved Pearson *r* = 0.222, a single attention block reached *r* = 0.779, and two blocks performed comparably (*r* = 0.772). LOH classification followed the same pattern (Fig. 2c): AUC rose from 0.683 at baseline to 0.943 with one attention block and 0.944 with two. The single-block improvement with plateau at two blocks indicates that the informative inter-segment relationships are captured at a single attention layer, consistent with CNAs as pairwise or local co-alterations rather than higher-order dependencies.

To test cross-platform generalisation we evaluated the TCGA-trained CNA models on MSK-CHORD^28^ panel sequencing. Panel-filtered segments were quantile-normalised onto the TCGA training distribution before inference to reconcile differences in copy-number calling and genomic coverage; because panels lack allele-specific copy-number calls, we used a *NoLOH* variant trained without the LOH head. Across the WES-to-panel shift (Fig. 2d), one attention block achieved *r* = 0.727 on MSK-CHORD and two blocks *r* = 0.746, whereas the attention-free baseline dropped to *r* = 0.053. Inter-segment copy-number patterns learned from whole-exome data therefore transfer to the limited genomic coverage of targeted panels.

### Pan-cancer tumour-type classification

We next asked whether these per-modality representations are sufficient to classify tumour types. Per-patient features were obtained by concatenating, for each modality, mean and max pools of the per-token embeddings together with log-transformed variant and segment counts. This per-patient *genomic fingerprint* (Fig. 3a) serves as input to a multi-layer perceptron classifier and is reused, unchanged, throughout the downstream analyses in this paper. Classification was performed across 23 TCGA cancer types (≥100 samples) using macro-fold nested cross-validation with 5 outer folds and 10 inner folds.

**Figure 3.**
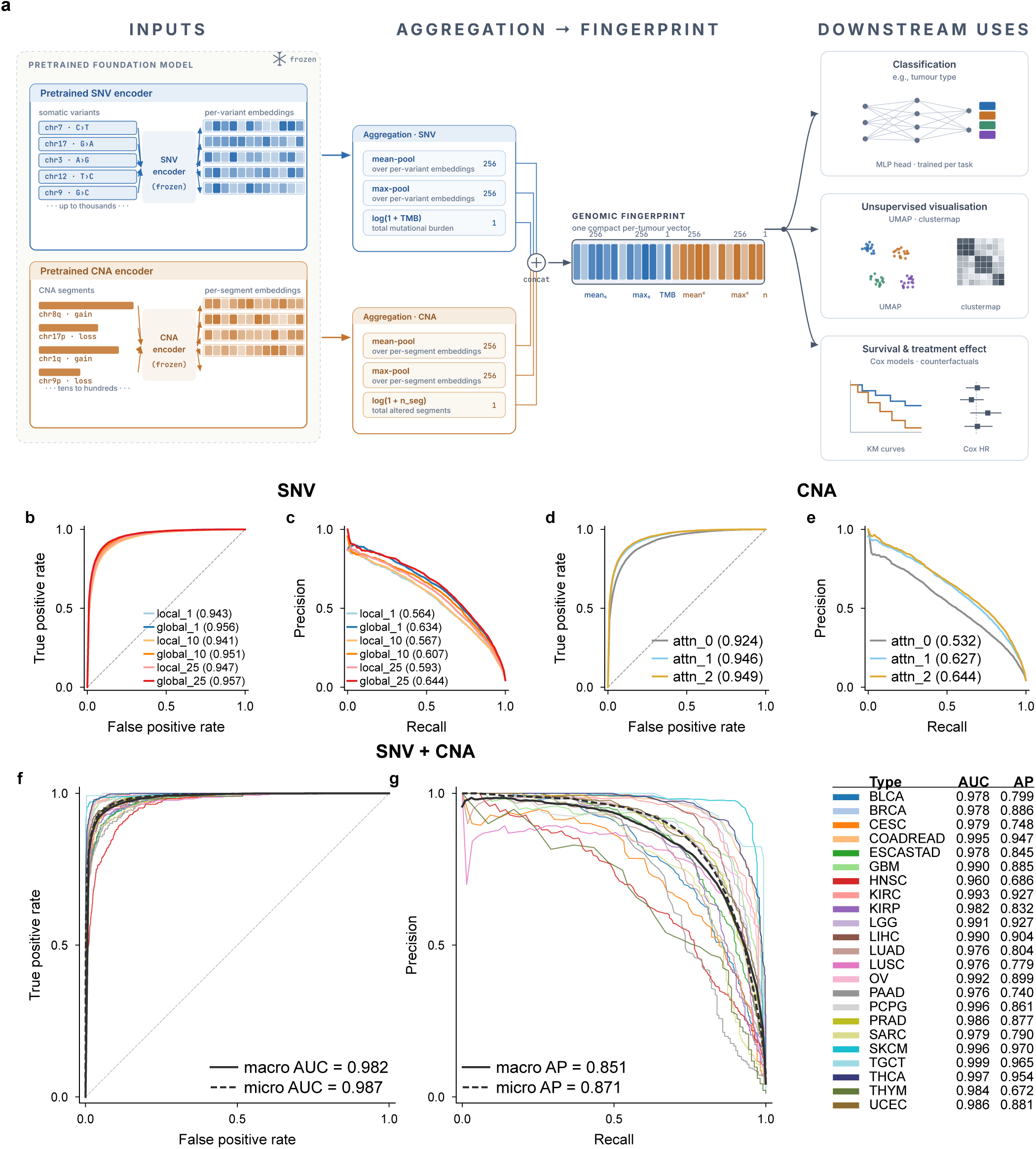
Tumour type classification using learned variant and copy-number representations. Multi-class tumour type classification across 23 TCGA cancer types (≥100 samples per type), evaluated using macro-fold nested cross-validation (5 outer folds, 10 inner folds). **a,** Schematic of the genomic-fingerprint construction. Per-variant and per-segment embeddings from the pretrained SNV and CNA encoders (Fig. 1a and Fig. 2a) are aggregated to per-patient features by concatenating per-modality mean and max pools with log-transformed variant and segment counts. The resulting fingerprint is reused as input across the downstream applications described in this paper: supervised classification (e.g., tumour type), unsupervised visualisation (UMAP, clustermaps), and survival/treatment-effect modelling. **b,** Macro-average ROC curves for SNV-based tumour type classification. Models compared: local attention only (local_1, local_10, local_25 with 1bp, 10bp, 25bp flanking context) and global attention (global_1, global_10, global_25). AUC values indicated in legend. **c,** Macro-average precision-recall curves for SNV-based classification. Same models as panel (b). Average precision (AP) values indicated in legend. **d,** Macro-average ROC curves for CNA-based tumour type classification. Models compared: baseline with no self-attention (attn_0) and models with 1 and 2 self-attention blocks (attn_1, attn_2). AUC values indicated in legend. **e,** Macro-average precision-recall curves for CNA-based classification. Same models as panel (d). AP values indicated in legend. **f,** Per-class ROC curves for combined SNV+CNA multimodal ensemble (global_25 SNV model + attn_2 CNA model). Individual curves shown for each of 23 tumour types. Micro-average and macro-average ROC curves shown as dashed lines. AUC values for all classes and averages indicated in legend. **g,** Per-class precision-recall curves for the SNV+CNA ensemble. Same model as panel (f). Random baseline shown as dashed horizontal line. AP values for all classes and averages indicated in legend.

Global-attention SNV models consistently outperformed local-only models (Fig. 3b–c): global_25 reached 0.644 macro-average precision (AP) versus 0.593 for local_25, with analogous gains at other context sizes. For CNA (Fig. 3d–e), one attention block (attn_1) reached 0.627 macro-AP and two blocks (attn_2) 0.644, compared with 0.532 for the attention-free baseline, a 21% relative improvement. Per-class ROC, PR, and confusion matrices for all SNV and CNA configurations are provided in Supplementary Figures 3–8.

Combining the two modalities substantially outperformed either alone (Fig. 3f–g): the multimodal ensemble (global_25 SNV + attn_2 CNA) achieved 0.851 macro-AP (0.871 micro-AP) and 0.982 macro-AUC (0.987 micro-AUC), compared with 0.644 macro-AP for the best single-modality model, indicating that somatic variants and copy-number alterations carry complementary information for tumour classification.

### Joint InfoNCE alignment

The independent SNV and CNA encoders used to produce the genomic fingerprint have no explicit pressure to agree about which tumour a sample comes from, since each was pretrained only on its own reconstruction task. We therefore jointly pretrained the two encoders with a bidirectional InfoNCE contrastive loss added to their reconstruction objectives (Fig. 4a): modality-specific projections of per-token backbone features are mean-pooled to sample-level embeddings, and InfoNCE treats same-sample SNV and CNA embeddings as positive pairs while pushing apart all other within-batch pairings, so that each encoder continues to learn modality-specific structure through its reconstruction loss while also producing features that agree on sample identity across modalities.

**Figure 4.**
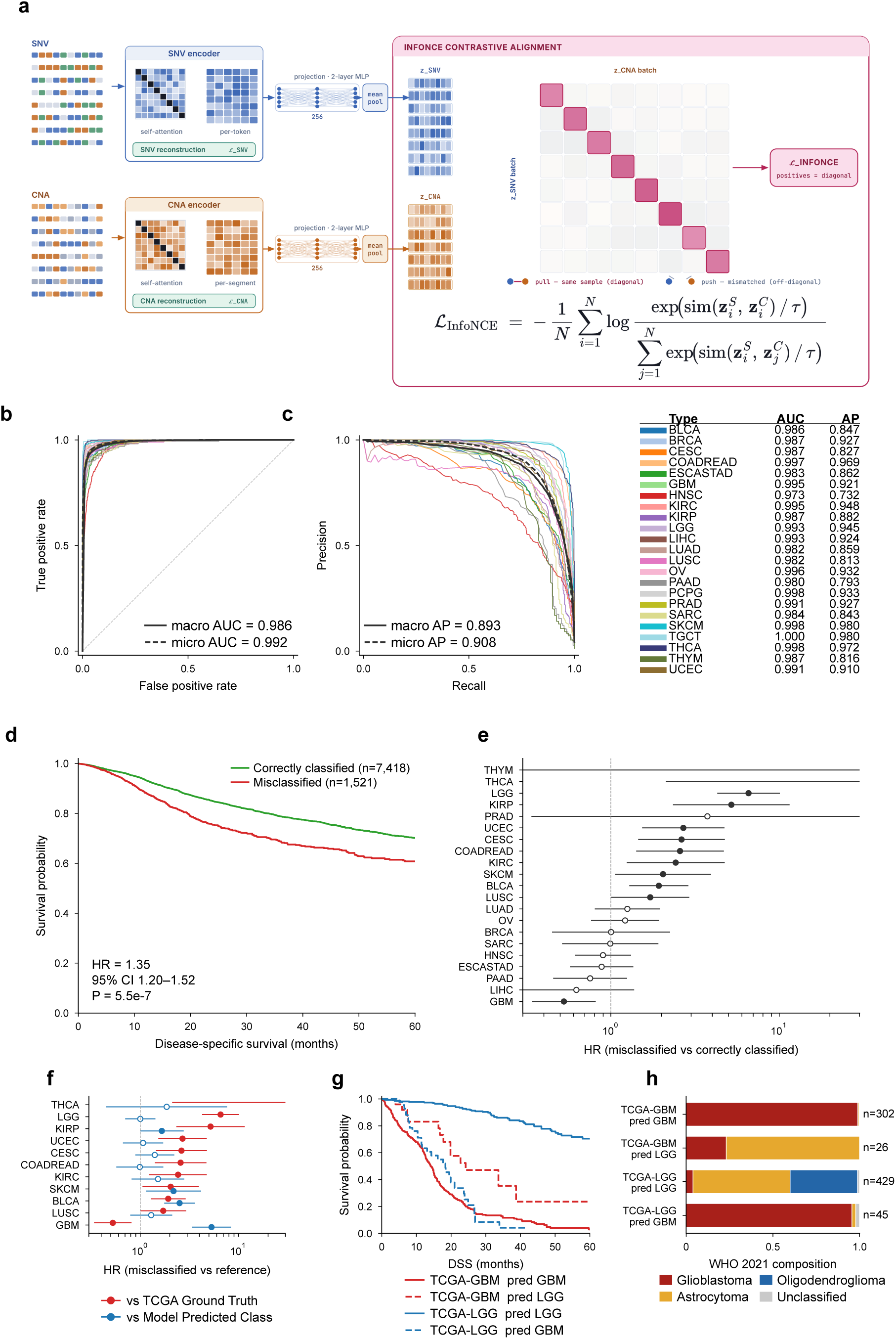
Joint SNV+CNA pretraining with InfoNCE and classifier concordance with disease-specific survival. **a,** Schematic of TESSERA, the joint SNV+CNA foundation model. The SNV and CNA encoders (Figs. 1a, 2a) are trained jointly on samples with both modalities; each retains its masked-reconstruction objective. Modality-specific projection heads map per-token backbone features to sample-level embeddings, over which a bidirectional InfoNCE contrastive loss treats same-sample SNV/CNA embeddings as positive pairs and all other within-batch pairings as negatives. **b–c,** Per-class ROC (b) and precision-recall (c) curves for tumour-type classification using the InfoNCE-aligned SNV+CNA ensemble, evaluated with the macro-fold nested cross-validation protocol of Fig. 3 across 23 tumour types. Micro- and macro-averages shown as dashed lines; AUC and AP values indicated in legends. **d,** Pan-cancer Kaplan–Meier curves of disease-specific survival (DSS) in correctly classified (green) versus misclassified (red) tumours across 8,939 TCGA samples with out-of-fold predictions, 60-month administrative censoring. The annotated hazard ratio is from a Cox model stratified by pathology class. **e,** Per-class forest of Cox hazard ratios (misclassified vs correctly classified) with 95% Wald CIs, one row per pathology class with ≥ 20 samples per arm. Filled circles denote *P <* 0.05; dashed vertical line at HR = 1; overflow whiskers truncated. **f,** Per-class paired forest restricted to the 11 classes with *P <* 0.05 in panel (e). Red (vs TCGA Ground Truth): HR of misclassified-*X* versus correctly-classified peers of the same TCGA pathology label (within-class Cox). Blue (vs Model Predicted Class): HR of misclassified-*X* versus correctly-classified peers of the classifier’s predicted class, stratified by predicted class. Filled circles denote *P <* 0.05 (HR significantly different from 1); hollow circles denote *P* 0.05, so a hollow blue circle indicates that misclassified samples’ survival is statistically indistinguishable from the classifier’s predicted class. **g,** Glioma case study, four-curve DSS Kaplan–Meier within TCGA GBM+LGG. Colour encodes TCGA pathology label (GBM red; LGG blue) and linestyle encodes classifier agreement (solid = correctly classified; dashed = misclassified). **h,** WHO 2021 label composition of the four Kaplan–Meier groups in (g), as stacked bars (oligodendroglioma orange; glioblastoma red; astrocytoma blue; unclassified grey).

The InfoNCE-aligned multimodal ensemble improved over the independently-pretrained concatenation baseline (Fig. 3f–g): it reached 0.986 macro-AUC and 0.992 micro-AUC on ROC analysis (Fig. 4b) and 0.893 macro-AP and 0.908 micro-AP on precision-recall analysis (Fig. 4c), a 0.042 macro-AP gain over the 0.851 concatenation baseline. This gain indicates that explicit cross-modal alignment during pretraining produces representations that better exploit the complementary information carried by somatic variants and copy-number alterations.

### Pathology-discordant tumours track genome-derived survival

With the InfoNCE-aligned tumour-type classifier (Fig. 4b–c) producing accurate calls, we asked what its occasional disagreements with the pathology-assigned label mean clinically. Using out-of-fold classifier predictions on 8,939 TCGA samples merged with disease-specific survival (DSS) from the Pan-Cancer Clinical Data Resource^29^ and 60-month administrative censoring, we compared misclassified to correctly-classified tumours in a pan-cancer Cox model stratified by pathology class (to control for class-specific baseline prognosis). Misclassified samples carried worse DSS than their pathology-matched correctly-classified peers (HR = 1.35, 95% CI 1.20–1.52; *P* = 5.5 × 10^−7^; Fig. 4d). Of 21 pathology classes with ≥ 20 samples in each arm, 11 reached *P <* 0.05 on a per-class Cox (Fig. 4e); LGG, KIRP, UCEC, CESC, COADREAD and BLCA had HRs ranging from 1.9 to 6.6, and GBM uniquely showed the opposite direction (HR = 0.52; *P* = 3 × 10^−3^), with misclassified GBMs surviving longer than their pathology peers. Per-class Kaplan–Meier curves are provided in Supplementary Figure 9.

Within these 11 classes we asked whether misclassified tumours track the survival of their classifier-predicted class rather than their pathology label. For each class *X* we computed two Cox HRs: misclassified-*X* versus correctly-classified peers of the same pathology class (`HR_actual`) and versus correctly-classified peers of the classifier’s predicted class (`HR_pred`; Fig. 4f). For 7 of 11 classes, `HR_actual` was different from 1 while `HR_pred` collapsed to 1 (*P >* 0.05). LGG illustrates the pattern: misclassified LGGs had `HR_actual` = 6.6 (*P <* 10^−17^) against correctly-classified LGG peers but `HR_pred` = 1.00 (*P* = 0.99) against correctly-classified peers of the classes the model assigned them, so their survival is captured by the classifier’s prediction, not by the LGG pathology label.

In glioma an external biological benchmark is available: the 2021 WHO revision reclassified IDH-mutant astrocytomas out of GBM into the LGG-family category, so WHO 2021 labels provide an independent reference against which the classifier’s pathology-disagreements can be tested.^30^ Within the TCGA-labelled LGG cohort, samples misclassified as GBM had DSS indistinguishable from correctly-classified GBMs (`HR_pred` = 0.83; *P* = 0.34; Fig. 4g) and were enriched for WHO 2021 glioblastoma (Fig. 4h), so the classifier identified IDH-WT tumours mislabelled as LGG under older criteria. TCGA-labelled GBMs misclassified as LGG were correspondingly enriched for WHO 2021 astrocytoma, the IDH-mutant entity reclassified out of GBM in the 2021 scheme, and showed intermediate DSS (Fisher’s exact *P* = 8 × 10^−24^ within TCGA-GBM). The classifier’s disagreements with pathology in both directions recapitulate the genomic reclassification subsequently codified by the WHO; the genomic fingerprint carries prognostic information beyond what the pathology label encodes.

### Unsupervised prognostic biomarkers

We next asked whether the representation also captures clinical structure *within* a tumour type. For each of four cohorts we projected the joint SNV+CNA features into two dimensions with UMAP, then asked (i) in tumour types with established discrete molecular subtypes, whether those subtypes are recovered as separable regions of the unsupervised manifold, and (ii) in tumour types whose underlying biology is more continuous, whether the genomic manifold itself encodes a continuous risk gradient. Survival was TCGA disease-specific survival (DSS)^29^ truncated at five years.

In cohorts with established molecular subtypes, the unsupervised projection recovered the known subtype structure without access to any subtype labels. In glioma (GBM + LGG, n = 865; Fig. 5a), the three WHO 2021 primary classes (astrocytoma, glioblastoma, oligodendroglioma)^30^ separated into distinct manifold regions, consistent with the classifier-level finding in Fig. 4g that TESSERA aligns with the 2021 WHO reclassification rather than the legacy GBM/LGG histologic labels. In endometrial carcinoma (UCEC, n = 458; Fig. 5b), the MSI-hypermutated and TP53-mutant copy-number-high serous-like subtypes^31^ occupied distinct regions of the UMAP. The cohort carries two copy-number-low endometrioid labels: one from the original TCGA 2013 classification,^31^ and one from the PanCancer Atlas reannotation.^9^ These overlapped on the projection, consistent with their shared biology: a copy-number-quiet, PIK3CA/PTEN/PIK3R1-driven entity.^31^

**Figure 5.**
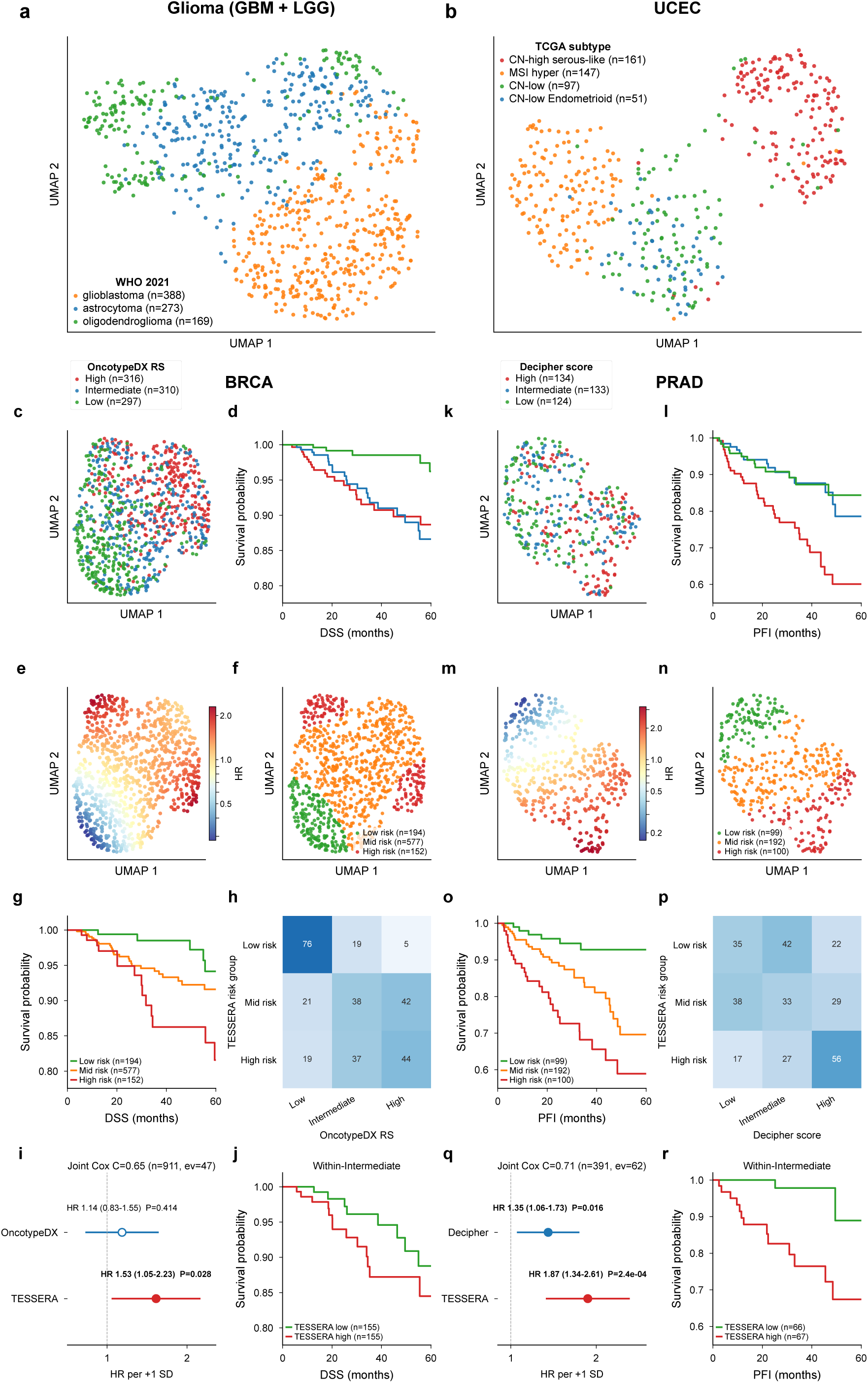
Unsupervised stratification of tumour cohorts from joint SNV+CNA features. Per-sample joint SNV+CNA embeddings from TESSERA were aggregated (mean + max pool), scaled, and projected to two dimensions with UMAP. **a,** Glioma (GBM + LGG, n = 865) coloured by WHO 2021 primary class. **b,** Endometrial carcinoma (UCEC, n = 458) coloured by TCGA molecular subtype. **c,** Breast cancer (BRCA, n = 933) coloured by OncotypeDX RS tertile. **d,** Kaplan–Meier curves for BRCA by OncotypeDX RS tertile. **e,** Same BRCA UMAP coloured by continuous TESSERA risk score (warmer = higher hazard). **f,** Same BRCA UMAP coloured by three TESSERA risk groups (low, intermediate, high) from the continuous score in (e). **g,** Kaplan–Meier curves for BRCA by TESSERA risk group. **h,** Concordance heatmap: percent of each TESSERA risk group (rows) falling in each OncotypeDX RS tertile (columns). **i,** Joint Cox forest for BRCA: hazard ratio per +1 SD of OncotypeDX RS and of TESSERA risk, with both predictors entered together. Markers, hazard ratio; whiskers, 95% Wald CI; dashed line, HR = 1; joint C-index in panel title. **j,** Kaplan–Meier curves for BRCA patients in the OncotypeDX-Intermediate stratum, split at the median TESSERA risk. **k–r,** As (c–j) for prostate adenocarcinoma (PRAD, n = 391), with the 22-gene Decipher score replacing OncotypeDX RS as the categorical and continuous comparator.

Breast cancer (BRCA, n = 933; Fig. 5c–j) and prostate adenocarcinoma (PRAD, n = 391; Fig. 5k–r) showed a different organisation: samples occupied a continuous genomic manifold along which tumours varied smoothly (Fig. 5c,k). For these two cohorts we used the field-standard transcriptomic prognostic biomarker as the categorical comparator, tertile-binned within the cohort: in BRCA the OncotypeDX 21-gene Recurrence Score (RS), reconstructed from TCGA RNA-seq via the Paik 2004 formula^32^ as released in the DLRS resource;^33^ in PRAD the 22-gene Decipher metastasis classifier,^34,35^ taken from the curatedPCaData uniform processing of TCGA-PRAD.^36^ The resulting Low/Intermediate/High strata imposed only partial structure on the genomic manifold (Fig. 5c,k) and produced significant but modest survival separation (Fig. 5d,l), consistent with a continuous rather than discrete underlying biology in these two tumour types.

Because the BRCA and PRAD genomic profiles populated a continuous manifold rather than discrete clusters, we fit a degree-1 polynomial Cox proportional-hazards model with interaction on the two UMAP coordinates to read a per-sample risk score directly off the manifold geometry. The fitted score varied smoothly across the projection (Fig. 5e,m), and partitioning it into three groups (Fig. 5f,n) yielded significant survival stratification in both cohorts (Fig. 5g,o). The model-derived strata only partially recapitulated the published transcriptomic comparator (Fig. 5h,p): the high-risk stratum included sizeable fractions of all three comparator tertiles. This discordance identifies patients in the *favourable* comparator strata whose joint SNV+CNA profile places them in a high-risk region of the manifold. To test whether the TESSERA risk score adds information beyond the established transcriptomic biomarker, we fit a joint Cox model with both predictors as standardised continuous covariates (Fig. 5i,q). In BRCA, the TESSERA score remained significant (HR_TESSERA_ = 1.53, 95% CI 1.05–2.23, *P* = 0.028; DSS, *n* = 911, 47 events, joint *C* = 0.65, Spearman *ρ* = 0.49) while OncotypeDX RS collapsed to null after mutual adjustment (HR_OncotypeDX_ = 1.14, 95% CI 0.83–1.55, *P* = 0.41), reflecting that the genomic information shared by the two correlated scores is captured at least as well by TESSERA. In PRAD, both predictors retained significance and the TESSERA score was the stronger (HR_TESSERA_ = 1.87, 95% CI 1.34–2.61, *P* = 2.4 × 10^−4^; HR_Decipher_ = 1.35, 95% CI 1.06–1.73, *P* = 0.016; PFI, *n* = 391, 62 events, joint *C* = 0.71, *ρ* = 0.31), reflecting complementary information. Within the OncotypeDX-Intermediate (BRCA) and Decipher-Intermediate (PRAD) strata, a TESSERA median split further stratified outcomes (Fig. 5j,r), so TESSERA adds resolution where the clinical risk score is least decisive. The TESSERA score is therefore independently prognostic of the established transcriptomic comparator in both cohorts, with two patterns: in BRCA the comparator’s signal is largely subsumed, whereas in PRAD the two scores carry orthogonal information.

### Counterfactual chemotherapy biomarkers

Oncology biomarkers are classified as prognostic, informing outcome regardless of therapy, or predictive, identifying patients who benefit differentially from one treatment over another; predictive biomarkers are rarer and more directly actionable.^27^ We applied TESSERA to two first-line metastatic chemotherapy settings in which clinical equipoise has persisted for two decades without a validated biomarker: FOLFOX versus FOLFIRI in colorectal cancer (CRC; n = 1,204 and 248 after pipeline-eligibility exclusions) and FOLFIRINOX (FFX) versus gemcitabine plus nab-paclitaxel (GA) in pancreatic ductal adenocarcinoma (PDAC; n = 479 and 292). Randomised head-to-head trials have established survival equivalence between the CRC doublets,^37^ and current ESMO and NCCN guidelines list both as acceptable first-line backbones with selection guided by toxicity and patient factors rather than tumour biology.^38^ In PDAC, FOLFIRINOX^39^ and gemcitabine plus nab-paclitaxel^40^ were established as standards of care in separate pivotal trials without direct head-to-head comparison, and choice in practice is determined by performance status and comorbidity.

The pipeline is summarised in Fig. 6a. For each cohort, the joint genomic fingerprint was passed through a doubly-robust learner^41,42^ on PFS, producing a per-patient predicted treatment effect 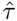 (arm-1 minus arm-0). An indifference threshold 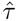_0_ partitioned patients into predicted-arm-1-favoured (FOLFOX in CRC; FOLFIRINOX in PDAC) and predicted-arm-0-favoured (FOLFIRI; gemcitabine plus nab-paclitaxel) strata, where 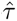_0_ is the value at which the cohort-level treatment-by-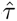 interaction Cox predicts no effect. All 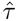 scores are out-of-fold under 10-fold outer / 5-fold inner stratified nested cross-validation.

**Figure 6.**
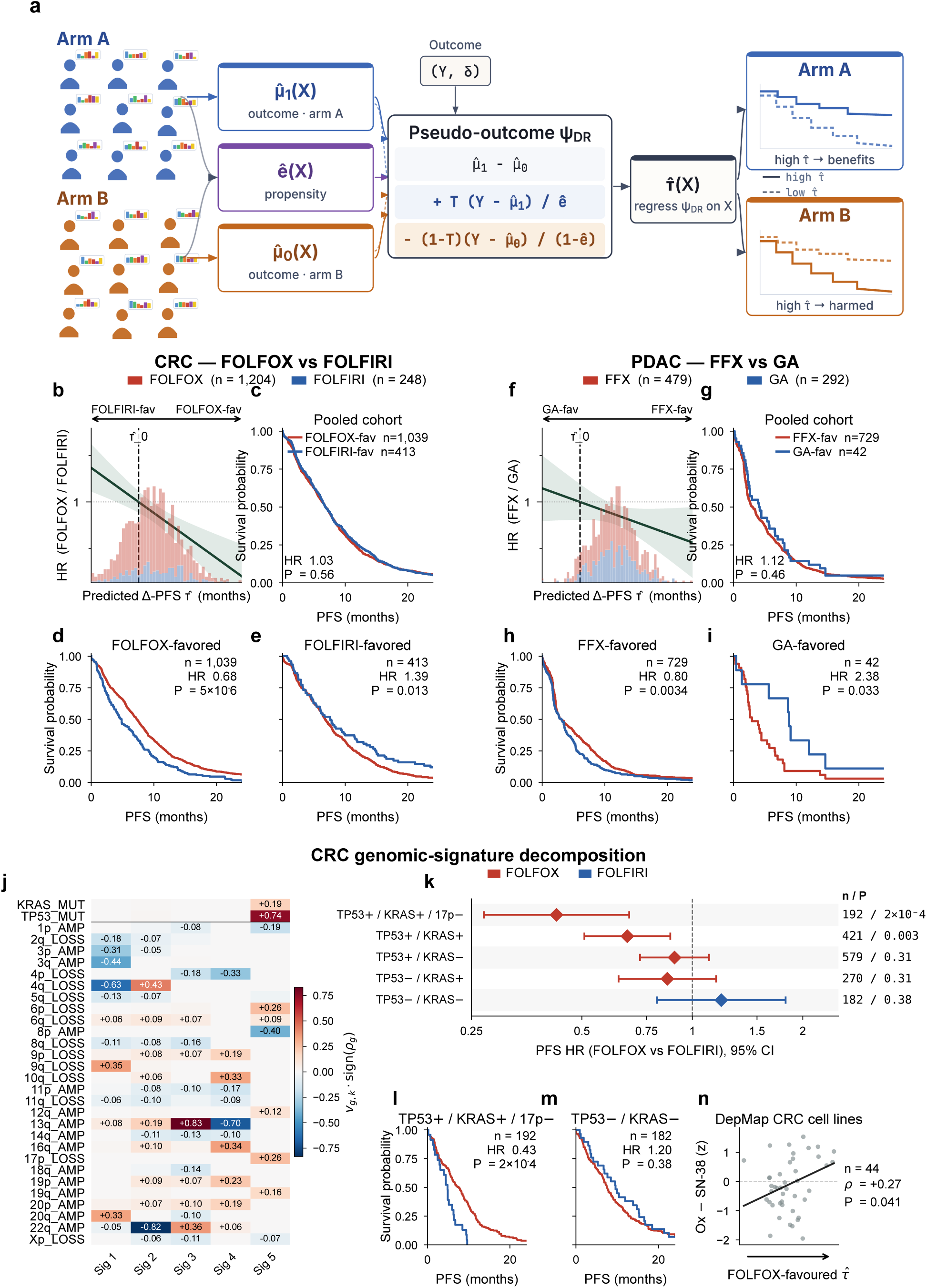
Doubly-robust counterfactual treatment-effect pipeline applied to first-line metastatic CRC and PDAC. TESSERA model (Fig. 4a) applied without retraining to two MSK-CHORD cohorts: metastatic CRC, FOLFOX (n = 1,204) versus FOLFIRI (n = 248) in panels b–e; metastatic PDAC, FOLFIRINOX (n = 479) versus gemcitabine plus nab-paclitaxel (n = 292) in panels f–i. Arm-1 treatments are red, arm-0 blue. All Kaplan–Meier curves show progression-free survival truncated at 24 months for legibility, with administrative censoring at 36 months. **a,** Schematic of the DR-learner pipeline. Per-arm Cox restricted-mean predictions 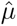_1_(*X*) and 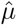_0_(*X*) and a propensity model *ê*(*X*) enter the doubly-robust pseudo-outcome *ψ*_DR_(*X*), on which a sparse partial-least-squares fit 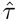(*X*) produces a per-patient predicted Δ-PFS. The interaction-Cox indifference threshold 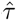_0_ separates predicted-arm-1-favoured (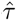 *>* 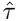_0_) from predicted-arm-0-favoured (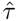 *<* 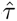_0_) patients. **b,** CRC. Per-arm 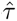 histograms with the analytic interaction-Cox HR(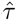) curve and shaded 95% Wald CI overlaid (left axis, log scale). Dashed line marks 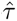_0_; arrows above the panel indicate the two predicted-favoured regions. **c,** CRC pooled-cohort PFS, regardless of arm, split at 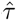_0_. **d, e,** CRC arm-stratified PFS within the predicted-FOLFOX-favoured stratum (d) and the predicted-FOLFIRI-favoured stratum (e). **f–i,** PDAC analogues of panels b–e. **j,** Heatmap of the *K* = 5 penalised matrix decomposition of the per-(patient, feature) attribution matrix toward CRC 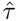. Rows: 30 features (2 SNV genes, 28 chromosomal arms), each a top-eight loader of at least one signature. Columns: signatures Sig 1–5; each cell shows that feature’s loading on that signature. Cell colour is the directional loading *v_g,k_* sign(*ρ_g_*), where *ρ_g_* is the Spearman correlation between feature *g*’s attribution column and its raw biological observable; row labels (AMP/LOSS for arms, MUT for SNVs) encode the same direction so that red uniformly indicates a positive load on the FOLFOX axis. **k,** Forest plot of within-subgroup CRC PFS Cox HR (95% Wald CI) for five exclusive TP53/KRAS/17p genotypes, ordered from triple-altered (TP53^+^/KRAS^+^/17p^−^) to double-wildtype. Right-hand column: within-subgroup *n* and Wald *P*. **l, m,** CRC arm-stratified PFS within the TP53^+^/KRAS^+^/17p^−^ triple-altered subgroup (l) and the TP53^−^/KRAS^−^ double-wildtype subgroup (m). **n,** DepMap CRC cell-line validation. MSK-CHORD-trained 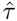 on CCLE/DepMap colorectal cell lines (x-axis) versus z-scored CTRPv2 sensitivity preference for oxaliplatin (FOLFOX backbone) over SN-38 (FOLFIRI active metabolite) (y-axis). Black line: linear regression. Right-pointing arrow: direction of increasing 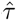. *n*, Spearman *ρ*, and one-sided *P* annotated. In each KM panel (c–e, g–i, l–m), *n* is the within-stratum sample size, HR is the Cox estimate of arm 1 versus arm 0, and *P* is the two-sided Wald *P*.

In CRC, the threshold partitioned the cohort into 1,039 predicted-FOLFOX-favoured and 413 predicted-FOLFIRI-favoured patients (Fig. 6b). The pooled split by 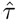_0_ produced overlapping PFS curves (HR = 1.03, *P* = 0.56; Fig. 6c), so the score carries negligible prognostic information and any predictive signal should emerge only under arm stratification. Within the FOLFOX-favoured stratum, FOLFOX outperformed FOLFIRI on PFS (HR = 0.68, 95% CI 0.58–0.80; *P* = 5 × 10^−6^; Fig. 6d); within the FOLFIRI-favoured stratum the direction reversed and FOLFIRI outperformed FOLFOX (HR = 1.39, 95% CI 1.07–1.81; *P* = 0.013; Fig. 6e). The two strata sit on opposite sides of the null in the same cohort under the same pipeline, the direction-reversal that defines a predictive biomarker. The same reversal held on overall survival, an endpoint not used to train 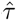: FOLFOX-favoured HR = 0.77 (0.63–0.95; *P* = 0.016) and FOLFIRI-favoured HR = 1.42 (1.03–1.97; *P* = 0.033); per-stratum OS Kaplan–Meier curves are in Supplementary Figure 10. Applied to the SEER-Medicare first-line stage IV CRC distribution^43^ (79.3% FOLFOX, 20.7% FOLFIRI, no measured OS difference between arms), and assuming biomarker classification is independent of current treatment assignment (consistent with selection by toxicity rather than tumour biology), the biomarker would reassign approximately 37% of patients: 22.6% of the cohort would move from FOLFOX to FOLFIRI, gaining a within-stratum median +0.9 months PFS and +6.9 months OS, and a further 14.8% would move from FOLFIRI to FOLFOX, gaining +2.4 months PFS and +8.4 months OS. These OS gains are on the same scale as the established benefit of adding biologics to first-line stage IV CRC chemotherapy.^44,45^ To localise the predictive signal within the representation, we re-ran the DR-learner on three nested feature subsets: local-attention only, local + global-attention, and the full local + global + InfoNCE-aligned features. The strata-level direction-reversal sharpened with each tier, with the largest contributions from the within-modality global-attention and cross-modal InfoNCE-aligned features (Supplementary Figure 11).

Applied unchanged to the PDAC cohort (Fig. 6f–i), the same pipeline produced an analogous bilaterally direction-reversed PFS signature, scaled to PDAC’s stronger overall preference for FOLFIRINOX. The indifference threshold partitioned the cohort into 729 predicted-FOLFIRINOX-favoured patients and 42 predicted-gemcitabine-plus-nab-paclitaxel-favoured patients, an exceptional-responder minority of ∼ 5% in whom the model predicts the less-aggressive arm. The pooled-cohort split by 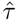_0_ was non-prognostic (HR = 1.12, *P* = 0.46; Fig. 6g), and the within-stratum arm comparisons direction-reversed: predicted-FOLFIRINOX-favoured patients fared better on FOLFIRINOX (HR = 0.80, 0.69–0.93; *P* = 0.003; Fig. 6h), and predicted-gemcitabine-plus-nab-paclitaxel-favoured patients fared better on that arm (HR = 2.38, 1.07–5.29; *P* = 0.033; Fig. 6i). OS validation was directionally consistent but did not reach significance, with FFX-favoured HR = 0.93 (0.79–1.09; *P* = 0.37) and GA-favoured HR = 1.27 (0.55–2.93; *P* = 0.58). Two factors plausibly contribute: the 42-patient GA-favoured stratum is event-poor, and PDAC’s aggressive natural history can dissociate PFS prediction from OS benefit, since rapid post-progression decline compresses survival differences that are visible on progression. The feature-slice ablation pattern recapitulated CRC’s (Supplementary Figure 11, PDAC panels): the direction-reversal sharpened with each tier and was largest under the full representation. In both cohorts, TESSERA produces a treatment-selection score satisfying the direction-reversal definition of a predictive biomarker, driven primarily by the global-attention and cross-modal InfoNCE-aligned features.

To localise the genomic basis of the CRC predictive direction, we composed the DR-learner’s post-aggregation chain into a single effective coefficient vector *β*_eff_, unwound the per-modality mean and max pools to obtain a per-(patient, variant or segment) attribution toward 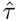, and aggregated to gene symbols for SNVs and to chromosomal arms for copy-number segments (the median MSK-IMPACT segment spans roughly 29 Mb, well above gene scale). A penalised matrix decomposition^46^ at rank *K* = 5 produced four signatures dominated by copy-number arm patterns and one combinatoric SNV/CNA signature whose top loadings were KRAS, TP53, and 17p loss (Fig. 6j, Sig 5). Reducing this signature to a binary genotype rule, FOLFOX-favoured if a patient carried both TP53 and KRAS coding mutations together with arm-level 17p loss (length-weighted log2 ratio *<* −0.2), separated the cohort into two genomic extremes with opposing arm preferences (Fig. 6k). TP53^+^/KRAS^+^/17p^−^ patients (*n* = 192) had a within-stratum PFS HR of 0.43 (95% CI 0.27–0.67; *P* = 2 × 10^−4^) favouring FOLFOX (Fig. 6l), and TP53^−^/KRAS^−^ double-wildtype patients (*n* = 182) had an HR of 1.20 (0.80–1.80; *P* = 0.38) trending toward FOLFIRI (Fig. 6m). The three intermediate genotypes (TP53-only, KRAS-only, and double-mutant with 17p intact) interpolated between the two endpoints (Fig. 6k). The rule is a coarsened, hand-readable approximation of one of the five DR-learner signatures, covering the genomic extremes (∼26% of the cohort) rather than reproducing the cohort-level bilateral direction reversal of the continuous 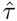 score; it expresses a dominant CRC predictive axis as three features already on every MSK-IMPACT report. The same decomposition applied to PDAC at rank 10 (Supplementary Figure 12) isolated two bilaterally direction-reversed signatures (Sig 8 and Sig 9; treatment-by-signature interaction *P* = 0.007 and 0.008). Sig 8 reduces to a three-feature rule: TP53 coding mutation, 17p intact (log2 ≥ −0.2), and 20q amplified (log2 *>* +0.2; substituted for the rarer 10q-amplification axis at the top of Sig 8). Patients matching the triplet (*n* = 61) had a within-stratum PFS HR of 0.53 (95% CI 0.30–0.92, *P* = 0.025), substantially below the cohort baseline of ∼ 0.83, while patients matching none (*n* = 261) sat at baseline (HR = 0.82, *P* = 0.12). The PDAC triplet is one-way: it identifies an FFX-enriched subgroup but does not flip direction toward GA, and the bilateral structure visible in the cohort-level 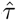 split (Fig. 6f–i, GA-favoured HR = 2.38) is not recoverable from any binary rule we tested. The two cohorts illustrate a single point: TESSERA’s predictive direction is interpretable enough to yield clinically tractable rules from features on standard sequencing reports, yet its full predictive resolution exceeds what those sparse rules capture.

To test whether 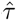 reflects tumour-intrinsic chemo-response biology or features specific to the MSK-CHORD population, we applied the frozen MSK-CHORD-trained 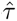 to colorectal cell lines from the Cancer Cell Line Encyclopedia / DepMap,^47^ an orthogonal cohort with no clinical context, no patient-level confounders, and drug response measured under uniform in-vitro conditions. Across the 44 CRC cell lines with both oxaliplatin and SN-38 dose-response measured in CTRPv2,^48^ 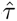 correlated with the FOLFOX-over-FOLFIRI sensitivity preference in the predicted direction (Spearman *ρ* = +0.27, one-sided *P* = 0.041; Fig. 6n). The same MSK-CHORD-trained score, applied without retraining and without any access to the in-vitro drug-response data, therefore recovers the FOLFOX-vs-FOLFIRI selection axis from intrinsic genomic features alone, indicating that the predictive signal reflects underlying tumour biology rather than artefacts of the training cohort.

## Discussion

We present TESSERA, a self-supervised foundation model for the cancer genome. A single representation, frozen at inference, supports variant pathogenicity prediction; pan-cancer tumour-type classification in which the model’s disagreements with pathology track class-specific prognosis better than the pathology label itself; unsupervised recovery of established molecular subtypes; prognostic risk stratification independent of clinically utilised transcriptomic scores; and counterfactual estimation of first-line chemotherapy benefit that satisfies the formal direction-reversal criterion of a predictive biomarker.

This work sits at the intersection of three existing threads in cancer machine learning: supervised pan-cancer tumour-type classifiers on clinical-sequencing features, including the clinically deployed GDD-ENS;^24,25^ supervised multi-modal pan-cancer survival models fusing genomic with pathology or expression features;^26^ and unsupervised somatic representations dominated by mutational-signature decompositions.^10^ Our contribution unifies these threads in one model: a self-supervised representation jointly trained on both somatic modalities and applied without retraining to a range of downstream oncology tasks.

For variant pathogenicity, the closest comparators are protein-centric foundation models that score missense effects (notably AlphaMissense^20^) and the genome-wide ensemble CADD;^49^ all are trained on germline variation or protein-family conservation and score each variant in isolation. TESSERA instead treats pathogenicity as a linear probe on a representation pretrained to reconstruct masked somatic alleles in tumour context, and uses masked-token reconstruction accuracy itself as a per-variant confidence filter, a cancer-specific affordance that germline or protein-only predictors do not offer.

The classifier-concordance analysis shows that misclassified tumours track the survival of the class the genome-derived classifier assigns them, not their pathology-assigned class. In glioma this is verified by external ground truth: misclassified cases align with WHO 2021 reassignments^30^ that were codified only after the original TCGA labels were applied. The unsupervised projections of glioma and endometrial carcinoma^31^ point the same way: the joint representation recovered their established discrete molecular subtypes without access to subtype labels, in line with a field that has moved from CMS in colorectal cancer^50^ to ProMisE adoption in endometrial carcinoma^51^ toward molecular taxonomies that the genome makes legible. Where external molecular ground truth exists, genome-derived classification therefore carries prognostic information beyond what the pathology label encodes. Breast and prostate cancer behaved differently: their genomes did not resolve into discrete subtypes but populated a continuous manifold, along which TESSERA read a per-sample risk score rather than recapitulating a known subtype scheme. This risk score was independently prognostic of the established transcriptomic biomarker (OncotypeDX^32^ in breast and Decipher^34,35^ in prostate), and in breast largely subsumed it.

The predictive-biomarker analyses operationalise a distinction well established but rarely cleanly isolated in the genomic-biomarker literature:^27,52^ a predictive biomarker requires a treatment-by-biomarker interaction, whereas a prognostic biomarker does not. The formal framework^53^ calls for this interaction to be either quantitative or, as in the canonical EGFR-gefitinib case,^54^ direction-reversing. Many published multi-gene oncology biomarkers are developed as single-arm prognostic scores,^32,34^ and the prognostic-versus-predictive distinction is rarely cleanly tested. Our joint-modality representation, fed through a doubly-robust counterfactual pipeline, produces a bilateral direction-reversing predictive biomarker in both the CRC FOLFOX-vs-FOLFIRI and PDAC FOLFIRINOX-vs-gemcitabine plus nab-paclitaxel comparisons on the PFS training endpoint, with a held-out OS validation that is significantly direction-reversed in CRC; in PDAC the OS signal was directionally concordant but not significant, plausibly reflecting both the small GA-favoured stratum and PDAC’s aggressive natural history that can dissociate PFS prediction from OS benefit (Supplementary Figure 10). The prognostic-versus-predictive decomposition^55^ should be the default lens for evaluating future genomic biomarkers on real-world clinicogenomic data.

The directly comparable prior work for the FOLFOX-vs-FOLFIRI predictive task is the FOLFOXai signature, an ensemble of five 1,000-tree random forests over a 67-gene panel of somatic mutations and copy-number calls reported by Abraham et al..^56^ Applied to the same MSK-CHORD first-line stage IV CRC cohort under faithful Abraham-paper conventions (five random-forest configurations, ensemble-mean *p*_IB_, 3% no-call buffer, time-to-next-treatment IB/DB labels at 270 days), the signature recovered a strong pooled prognostic effect on both endpoints (PFS HR = 0.79, 95% CI 0.69–0.91, *P* = 9 × 10^−4^; OS HR = 0.70, 95% CI 0.60–0.82, *P* = 1 × 10^−5^). However, neither arm-stratified comparison reached significance on either endpoint and no direction reversal was observed (within the predicted-IB stratum, FOLFOX vs FOLFIRI: PFS HR = 0.88, *P* = 0.16; OS HR = 0.94, *P* = 0.57; within the predicted-DB stratum: PFS HR = 0.86, *P* = 0.36; OS HR = 1.18, *P* = 0.39; Supplementary Figure 13). Under the Ballman framework, the FOLFOXai signature is therefore prognostic on this cohort but does not satisfy the direction-reversal criterion of a predictive biomarker. This follows from the training design. FOLFOXai is trained on a single arm, the FOLFOX cohort with IB/DB labels from 270-day time-to-next-treatment discontinuation (TTNTD), so its training signal is which patients fare well on FOLFOX rather than how outcomes differ between arms; single-arm training of this form is intrinsically prone to learning prognostic rather than predictive features, because the model cannot see the counterfactual outcome under the alternative treatment. TESSERA’s DR-learner, by contrast, trains on both arms simultaneously and targets the treatment-effect contrast directly via a doubly-robust pseudo-outcome that is consistent if either the per-arm outcome model or the propensity model is correctly specified, controlling for both prognostic and treatment-assignment confounding by construction. Two caveats accompany this analysis. Abraham’s 67-gene panel was originally trained on the broader Caris Molecular Intelligence platform; only 28 of the 67 genes (42%) are profiled by MSK-IMPACT, and the remaining 39 are absent from MSK-CHORD calls, so the comparison strictly tests the MSK-IMPACT-overlap portion of the FOLFOXai feature set rather than the full Abraham signature. Separately, Abraham’s discrete-class biomarker abstains in the 3% buffer around *p*_IB_ = 0.5, excluding 23% of patients (336 of 1,443) from any prediction; TESSERA’s continuous 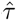 assigns a direction to every patient, a coverage property of the continuous-score architecture rather than a tuning choice.

Beyond satisfying this formal definition, if prospectively validated, the per-patient benefit projected for biomarker-guided treatment selection is on a clinically meaningful scale. Re-assigning the ∼37% of patients identified by the model in the SEER-Medicare-projected analysis would deliver a within-stratum median OS gain of 7–8 months on the recommended regimen, on the same scale as the established benefits of adding biologic therapy to chemotherapy in first-line metastatic CRC: ∼5–6 months OS for biology-selected anti-EGFR (panitumumab in RAS-wildtype patients in the PRIME trial^44^) and ∼4–5 months OS for unselected anti-VEGF (bevacizumab added to chemotherapy^45^).

TESSERA’s treatment-selection score is also interpretable: each patient’s predicted treatment effect can be traced back to the specific somatic alterations driving it. Grouping these per-feature attributions into sparse genomic signatures yielded clinically tractable genotype rules in both cancers: a TP53^+^/KRAS^+^/17p^−^ rule isolating the FOLFOX-favoured genomic tail in CRC (Fig. 6j–m) and a one-way TP53^+^/17p intact/20q^+^ FFX-enrichment rule in PDAC (Supplementary Figure 12). The same analysis also bounds what interpretability yields: PDAC’s bilateral two-way prediction visible in the full 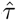 score is not recoverable from any sparse rule we tested, indicating that the model’s full predictive resolution lives in the joint genomic complexity rather than in any low-rank decomposition. These two findings are complementary. The interpretable rule validates that the model has learned biologically grounded patterns rather than spurious signal; the full continuous score, which captures the model’s complete predictive resolution, is what would be deployed clinically. Future work will probe the foundation model directly through its attention weights, which encode the variant-variant and segment-segment dependencies learned during pretraining and sit upstream of the post-aggregation linear step decomposed here.

The CRC predictive signal also survives transfer to a cohort with no clinical context: applied without retraining to 44 colorectal cell lines from CCLE/DepMap, the MSK-CHORD-trained 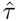 correlated with the FOLFOX-over-FOLFIRI sensitivity preference measured in CTRPv2 (Fig. 6n). Because the cell-line cohort shares none of MSK-CHORD’s patient-level confounders (no performance-status differences, no comorbidity profiles, no physician preferences), this orthogonal validation is consistent with the predictive signal reflecting tumour-intrinsic chemo-response biology rather than artefacts of the clinical training population.

Several caveats temper our conclusions, falling into three groups: pretraining-data scope, clinical-cohort design, and the standard of evidence. The pretraining corpus is the TCGA Pan-Cancer Atlas, which is dominated by treatment-naïve primary tumours and does not directly reflect the metastatic, post-treatment, and resistance states in which the predictive-biomarker applications are evaluated; the somatic modality is also scoped to SNVs, so small insertions and multi-nucleotide substitutions are not modelled as first-class events, a tokenisation-level scope decision rather than a structural limit. The clinical cohorts in turn come from routine care in MSK-CHORD rather than randomised trials, so treatment assignment is confounded by performance status, comorbidity, and physician preference, and standard caveats on observational causal inference apply throughout.^57,58^ Every clinical claim here is therefore retrospective: the standard for elevating a multi-gene biomarker from prognostic to predictive is prospective randomised validation,^59,60^ and we make no claim that any score here has met that bar.

The representation is directly computable on data already produced by routine panel-based NGS, so the translational next step is a prospective randomised evaluation of a model-derived treatment-selection score in first-line metastatic CRC or PDAC, following the TAILORx/MINDACT template. The most clinically actionable methodological extension, entirely unexplored here, is pretraining natively on large corpora of panel-based NGS, such as those catalogued in GENIE, rather than transferring from TCGA whole-exome data: a panel-native alternative would use roughly an order of magnitude more samples with greater metastatic and post-treatment representation, and would eliminate the WES-to-panel distribution shift. Other extensions include additional somatic modalities (gene fusions, structural variation, methylation, expression), whole-genome pretraining on PCAWG-scale data,^12^ cross-modal alignment between the genomic representation and a pathology foundation model, and end-to-end attention-MIL successors to the pool-then-Cox downstream pipeline.^61^ More broadly, the self-supervised framework over co-occurring genomic alterations is not specific to cancer and is in principle applicable to germline settings where polygenic variant co-occurrence underlies disease risk; demonstrations beyond the oncology context are deferred to future work.

A single self-supervised representation of the cancer genome encodes the structure of somatic alterations that is relevant for both biological discovery and clinical decision-making. It can serve variant interpretation, molecular typing, prognostic stratification, and predictive-biomarker discovery within one framework, in the way that ESM-2 serves protein sequence,^18^ Geneformer serves single-cell transcriptomics,^7^ and UNI serves computational pathology.^8^ With TESSERA, a foundation model is now also available for the cancer genome.

## Methods

### Data sources and cohorts

Training data come from the TCGA Pan-Cancer Atlas, with clinical outcomes for the TCGA analyses from the curated Pan-Cancer Clinical Data Resource;^29^ cross-platform validation for the SNV encoder uses GENIE; variant-pathogenicity labels come from ClinVar; cross-platform CNA validation and clinical-application cohorts were assembled from the MSK-CHORD release; and an orthogonal cell-line validation uses CCLE/DepMap with drug-response data from CTRPv2.

#### TCGA pan-cancer cohort

Somatic SNVs were obtained from the Multi-Centre Mutation Calling in Multiple Cancers (MC3) MAF file, comprising whole-exome sequencing (WES) from approximately 10,000 tumour samples. Variants were filtered to retain those with sequencing depth *>* 0, and variant allele frequency (VAF) was calculated as t_alt_count / (t_alt_count + t_ref_count). Long alleles (*>*10 nucleotides) were truncated to the first and last 5 nucleotides. Analysis was restricted to 31 solid tumour types. CNA segments were obtained from TCGA ABSOLUTE algorithm calls, which provide allele-specific copy-number estimates. Segment mean values (log2 copy-number ratio) were computed with tumour purity adjustment: observed copy number was calculated as purity × tumour_CN + (1 − purity) × 2, and segment mean was defined as log2(observed_CN / 2). Loss of heterozygosity (LOH) status was derived from allele-specific copy-number calls. SNV and CNA data were split at the patient level into training (75%) and validation (25%) sets using stratified sampling by tumour type, with the same patient-level assignments across modalities.

#### GENIE validation cohort

The Genomics Evidence Neoplasia Information Exchange (GENIE) consortium v18.0 public release was used as an independent panel-sequencing validation set for the SNV encoder. Each sample is annotated by sequencing assay (SEQ_ASSAY_ID) and tumour type (ONCOTREE_CODE). Mutation data were processed with the same filtering as TCGA (depth *>* 0, VAF as above). For evaluation, samples with more than 100 variants were excluded to reduce hyper-mutator influence on the cross-platform comparison.

#### ClinVar pathogenicity annotations

Variant pathogenicity labels for the pathogenicity-prediction task were obtained from ClinVar, a public archive of relationships between human variations and phenotypes, with variants labelled pathogenic or benign based on clinical interpretation and supporting evidence.

#### MSK-CHORD validation and clinical cohorts

The MSK-CHORD 2024 release was used both for CNA cross-platform validation and for the clinical treatment-effect analyses. CNA segments were obtained from MSK-IMPACT targeted panel sequencing across four panel versions (IMPACT341, IMPACT410, IMPACT468, and IMPACT505); samples sequenced with the IMPACT-HEME-400 panel were excluded to restrict analysis to solid tumours. For the cross-platform CNA validation, CNA segments were filtered and trimmed to the gene regions covered by each sample’s specific panel, using panel gene coordinates from the GENIE genomic-information files, with only the overlapping portions of each CNA segment retained; for the clinical treatment-effect analyses, the raw MSK-CHORD segments were used directly without panel trimming, both for computational tractability (per-gene panel explosion expands the per-sample segment count roughly tenfold, which is prohibitive for the full clinical cohorts under repeated nested cross-validation) and to match the coarser whole-exome segmentation scale the TCGA-pretrained CNA encoder was trained on. Both uses applied the quantile-normalisation step described below, passing MSK-CHORD segment-mean values onto the TCGA training distribution before inference. For the predictive-biomarker analyses, two clinical cohorts were assembled de novo from the MSK-CHORD per-patient regimen, line-of-therapy, stage, and outcome tables. The first cohort comprises stage-IV colorectal cancer patients receiving first-line FOLFOX or FOLFIRI (n = 1,699; 1,408 FOLFOX, 291 FOLFIRI); the second comprises stage-IV pancreatic ductal adenocarcinoma patients receiving first-line FOLFIRINOX or gemcitabine plus nab-paclitaxel (n = 889; 544 FOLFIRINOX, 345 gemcitabine plus nab-paclitaxel). Both cohorts are defined by a regimen positive list, so the triple-cytotoxic FOLFOXIRI regimen was excluded from the CRC cohort; concurrent bevacizumab and other biologic agents were allowed and not used to define the comparator arms. Progression-free survival was computed as the number of days from first-line treatment start (TREATMENT_START at `LINE_OF_THERAPY = 1`) to the MSK-CHORD-annotated first-progression date (`FIRST_PROGRESSION_DATE`); patients without a recorded `FIRST_PROGRESSION_DATE` were excluded from the predictive-biomarker analyses, and the remaining PFS times were administratively censored at the 36-month horizon used by the DR-learner second-stage regression (event flag zeroed for patients whose progression occurred past this horizon). After this exclusion the analytic cohorts comprised 1,452 CRC patients (1,204 FOLFOX, 248 FOLFIRI) and 771 PDAC patients (479 FOLFIRINOX, 292 gemcitabine plus nab-paclitaxel). Overall survival was measured from treatment start using `OS_MONTHS_FROM_TREATMENT_START`, with the event indicator derived from MSK-CHORD vital status (`OS_STATUS`) and administratively censored at 60 months (event flag zeroed for patients surviving past this horizon).

#### CCLE / DepMap and CTRPv2

Cancer cell-line somatic mutations and copy-number segments (with LoH annotations) were taken from the Cancer Cell Line Encyclopedia / DepMap 24Q2 release.^47^ Drug-response area-under-the-curve values for oxaliplatin and SN-38 (the FOLFOX backbone and the FOLFIRI active metabolite, respectively) were taken from the Cancer Therapeutics Response Portal v2 (CTRPv2).^48^ The intersection of cell lines with both drugs profiled and a colorectal lineage annotation defined the *n* = 44 CRC cell-line validation cohort.

### Model architecture

The model comprises two modality-specific encoders, an SNV encoder and a CNA encoder, and a joint configuration that couples them through modality-specific projection heads and a cross-modal InfoNCE objective. All encoders are built from multi-head attention blocks (12 heads, 12 dimensions per head) with GELU activations, pre-layer normalisation, and residual connections, followed by feed-forward layers (288 hidden dimensions). The SNV encoder uses three blocks each in its local-attention and global-attention modules; the CNA encoder uses zero, one, or two self-attention blocks across the configurations compared. All models were implemented in TensorFlow.

#### Input features and tokenisation

Reference genome sequences were obtained from NCBI (GRCh37, assembly GCF_000001405.25_GRCh37.p13) and extracted using pyfaidx. For the SNV encoder, each variant is represented by its reference allele, alternate allele, chromosome, genomic position, VAF, and flanking genomic context (1bp, 10bp, or 25bp on each side). Nucleotides are tokenised with a six-symbol vocabulary (A, C, G, T, deletion, unknown) and one-hot encoded. For the CNA encoder, each segment is represented by chromosome (one-hot encoded), normalised start and end coordinates, log-transformed segment length, segment mean (log2 copy-number ratio), and LOH status.

#### SNV encoder

The SNV encoder maintains two parallel embedding streams to enable masked prediction: a reference stream containing only genomic position information, and a mutation stream containing both position and allele information. This separation ensures that when predicting the alleles at a given position, the model cannot see those alleles directly and must infer them from context. The encoder has two attention stages. In the *local attention module*, each stream cross-attends to its flanking sequence context processed through 1D convolutional layers, with separate attention heads for 5’ and 3’ context; this stage learns sequence-dependent features such as trinucleotide mutational signatures. In the *global attention module*, the reference stream serves as the query and the mutation stream as key and value, with a diagonal mask preventing each variant from attending to itself; this stage learns sample-level mutational signatures and variant-variant co-occurrence. The reconstruction head projects the attended embeddings through dense layers (512 and 128 units, GELU) with separate output layers for reference and alternate allele prediction. We compared seven model configurations: (1) baseline with no sequence context; (2–4) local-only attention with 1bp, 10bp, and 25bp context; and (5–7) global attention (local + global) with 1bp, 10bp, and 25bp context.

#### CNA encoder

The CNA encoder also uses two parallel streams to prevent leakage during the segment-mean prediction task: a query embedding containing chromosome, start, end, and segment length, and a key/value embedding that additionally contains segment mean and LOH. The embedding layer projects concatenated features through three dense layers with GELU and layer normalisation. Inter-segment self-attention blocks then allow each segment to attend to every other segment in the sample, learning genome-wide patterns of copy-number co-alteration; a diagonal mask prevents self-attention. Prediction heads output segment mean (linear activation, regression) and LOH (sigmoid activation, binary classification). We compared configurations with 0, 1, and 2 self-attention blocks (attn_0, attn_1, attn_2). A *NoLOH* variant, trained with the LOH prediction head removed, is used throughout all cross-platform and clinical analyses where allele-specific copy-number calls are not available (panel-based cohorts).

#### Joint SNV+CNA model (TESSERA)

TESSERA combines the SNV and CNA encoders above and adds modality-specific projection heads for the cross-modal InfoNCE contrastive objective. Each projection head is a 2-layer MLP with 256 units per layer (one GELU-activated hidden layer and a linear output) applied to the per-token backbone features of its modality prior to any cross-modal interaction, with masked global average pooling to a sample-level embedding (project-then-mean). Pretraining details for all three configurations are given in the next section.

### Pretraining

#### Optimisation

All encoders were pretrained with the AdamW optimiser (*β*_1_ = 0.9, *β*_2_ = 0.97, weight decay 0.001), fixed learning rate 2 × 10^−4^, gradient clipping at max norm 10, and dropout rate 0.1. Training used early stopping on validation-set performance with patience 10.

#### Input subsampling

To enable efficient training while preserving informative alterations, SNVs were subsampled to a maximum of 1,000 variants per sample, prioritising commonly occurring mutations (present in ≥ 5 samples) so that recurrent variants were retained. CNA segments were subsampled to a maximum of 1,000 per sample, prioritising high-amplitude alterations using the score |Segment_Mean| + 0.5 × LOH.

#### SNV pretraining

The SNV encoder was pretrained on TCGA somatic variants under the parallel-stream architecture described above (*SNV encoder*): a reference stream that carries only genomic position and a mutation stream that carries position together with reference and alternate alleles. The reconstruction head predicts both alleles at every variant position from the attended embeddings; because the global-attention block uses the reference stream as query and the mutation stream as key and value with a diagonal self-attention mask, no token can observe its own alleles, and prediction must rely on the surrounding genomic context and on the pattern of other variants in the sample. Reconstruction performance was evaluated by per-variant accuracy for reference and alternate allele prediction.

#### CNA pretraining

The CNA encoder was pretrained with dual-task learning: continuous segment-mean regression and binary LOH classification. Segment-mean values (log2 copy-number ratios, purity-adjusted as described in Data sources) were passed in their native scale. Segment-mean prediction was evaluated by Pearson correlation and LOH classification by AUC-ROC.

#### Joint SNV+CNA pretraining with InfoNCE

The SNV and CNA encoders were combined and trained jointly on samples with both modalities. Each modality retained its native reconstruction objective (masked-token SNV, masked segment-mean + LOH CNA), and a bidirectional InfoNCE contrastive loss between the L2-normalised sample-level SNV and CNA embeddings was added with weighting coefficient 0.1. The InfoNCE loss was computed symmetrically (SNV→CNA and CNA→SNV) as temperature-scaled cross-entropy over the batch similarity matrix with the same-sample diagonal as the target, with the two directions averaged and temperature 0.1. Reconstruction losses were computed per-sample (averaging per-token losses within a sample before averaging across samples) so that samples with few tokens or segments contributed equally regardless of input size. A valid-pairs mask ensured that only samples with both modalities contributed to the contrastive loss; samples with only one modality still received their own reconstruction loss.

#### MSK-CHORD segment-mean normalisation

Because the TCGA-trained CNA encoders operate on raw segment-mean values, applying them to MSK-CHORD required reconciling the difference in segment-mean distributions between cohorts, which arises from differences in copy-number calling (TCGA ABSOLUTE with purity adjustment vs. MSK-IMPACT panel calls) and in genomic coverage (WES vs. targeted panels). Before inference, each MSK-CHORD segment-mean value was quantile-normalised onto the TCGA training distribution: MSK values were rank-mapped to the corresponding quantile of a reference vector formed by pooling TCGA training and validation segment means. This rank-based transform re-expresses MSK-CHORD segments at the TCGA amplitude scale while preserving within-sample relative ordering.

### Variant pathogenicity prediction

Variant embeddings were extracted from the pretrained SNV encoder and used for supervised pathogenicity prediction on ClinVar-annotated variants. Features were standardised and dimensionality was reduced using PCA (retaining 99.9% variance). A logistic regression classifier with balanced class weighting was trained to predict pathogenic vs. benign status. Two evaluation strategies were employed: (i) *unique variant* evaluation, where each unique variant (chr:pos:ref:alt) was assigned exclusively to training or test sets (75%/25% variant-level split), representing standard evaluation for novel variants; and (ii) *unique gene* evaluation, a more stringent protocol where test variants came from genes completely absent during training (75%/25% gene-level split), testing whether the model learns transferable sequence-level patterns rather than gene-specific associations. In both cases, we additionally evaluated the subset of variants where the SNV encoder had correctly predicted masked reference and alternate alleles during pretraining, testing whether masked-token reconstruction accuracy can serve as a per-variant confidence filter on the pathogenicity call. Performance was evaluated using ROC AUC, average precision (AP), and balanced accuracy.

### Tumour type classification

Learned embeddings from the pretrained encoders were aggregated to per-sample features for supervised tumour-type classification. SNV token-level embeddings were aggregated by concatenating mean and max pools plus the sample’s mutational burden (total variant count). CNA segment-level embeddings were aggregated analogously with segment count as an additional feature. Features were standardised with RobustScaler. A multi-layer perceptron (three hidden layers of 512, 256, and 128 units with batch normalisation, ReLU, and dropout 0.5; softmax output) was trained to predict tumour type. Models were optimised with Adam (default learning rate 10^−3^), batch size 32, up to 1,000 epochs with early stopping on validation loss (patience 25, best-weight restoration); variant and segment counts were log(1 + *x*)-transformed before concatenation with the per-modality mean and max pools. Classification was restricted to 23 TCGA cancer types with at least 100 samples; COAD and READ were merged into a single COADREAD class and ESCA and STAD into ESCASTAD, reflecting their near-indistinguishable somatic profiles and shared clinical classification.

To ensure unbiased performance estimates, we employed macro-fold nested cross-validation with 5 outer folds and 10 inner folds, so that each sample is tested exactly once with no leakage between training and evaluation. Multimodal analyses concatenated independently-pretrained global_25 SNV features and attn_2 CNA features on samples with both modalities available and used the same MLP and macro-fold nested CV. The same protocol was then applied to features from the InfoNCE-aligned joint model. Performance was reported using micro- and macro-average ROC AUC and average precision (AP).

### Classifier concordance and DSS

Out-of-fold predictions from the 5-outer-fold, 10-inner-fold macro-fold nested cross-validation of the InfoNCE-aligned SNV+CNA tumour-type classifier were merged with the curated TCGA Pan-Cancer Clinical Data Resource^29^ on `bcr_patient_barcode`. Disease-specific survival (DSS) was used as the endpoint, with 60-month administrative censoring (`time_i = min(time_i, 60 months)`; `event = 0` if `time_i > 60 months`) to match the predictive-biomarker analyses below. Ambiguous cause-of-death codes (`DSS_cr = 2`) were treated as censored.

#### Pan-cancer test

A Cox proportional-hazards model of the form DSS ∼ misclassified, stratified by pathology class, was fit on all samples. The stratification forces the misclassified-vs-correctly-classified comparison to happen within each pathology class and then aggregates the per-stratum partial likelihoods, controlling for class-specific baseline prognosis.

#### Per-class test

For each pathology class with at least 20 samples in both the correctly-classified and misclassified arms, an unstratified Cox model DSS ∼ misclassified was fit on that class alone, producing a class-specific HR with 95% Wald CI and P-value.

#### Misclassified-tracks-predicted test

For each class *X* with ≥ 20 misclassified and ≥ 20 correctly-classified samples: (i) HR_actual was computed from an unstratified Cox on (correctly-classified-*X* ∪ misclassified-*X*), (ii) HR_pred was computed from a Cox on (misclassified-*X* ∪ correctly-classified samples whose actual label matched the classifier-predicted label of any misclassified-*X* sample), stratified by the classifier’s predicted class, so that within each stratum the comparison is between misclassified-*X* samples routed to that predicted class and the correctly-classified peers of that predicted class.

#### Glioma case study

TCGA GBM and LGG samples were merged with the updated WHO 2021 reclassification table of Leiria et al.^30^ on `bcr_patient_barcode`, yielding 1,110 glioma samples annotated with a WHO 2021 label (glioblastoma, astrocytoma, oligodendroglioma, or unclassified). Of these, 865 with one of the three primary WHO 2021 classes (glioblastoma, astrocytoma, oligodendroglioma) enter the Fig. 5 unsupervised projection analysis; the unclassified subset is excluded only from that downstream analysis and retained here. A four-curve Kaplan–Meier compares the four (pathology label × classifier call) groups within the GBM+LGG cohort, and horizontal stacked bars show the WHO 2021 composition of each of the four KM groups. Fisher’s exact test on the 2 × 2 contingency (classifier call × WHO 2021 equal to glioblastoma vs. not) within TCGA-labelled GBM quantifies whether the classifier’s reassignments align with the WHO 2021 reclassification.

### Unsupervised prognostic biomarkers

For each of four cohorts (glioma GBM+LGG, n = 865, restricted to the three primary WHO 2021 classes *astrocytoma*, *glioblastoma*, *oligodendroglioma*; endometrial carcinoma UCEC, n = 458; breast cancer BRCA, n = 933; prostate adenocarcinoma PRAD, n = 391), per-sample joint SNV+CNA features from the InfoNCE-aligned encoder were aggregated to the sample level by concatenating per-modality mean and max pools, standardised with RobustScaler, and projected to two dimensions with UMAP (*n*_neighbors_ = 100, min_dist_ = 0.5, cosine metric); to stabilise the embedding against random-seed variation, ten independent UMAP fits were Procrustes-aligned to a reference seed and averaged to produce the consensus embedding used for downstream risk-score derivation. Sample sizes decrease as analyses add data dependencies: for BRCA, n = 933 carry valid joint embeddings, 923 additionally have DSS and OncotypeDX RS, and n = 911 enter the joint Cox after excluding records with zero follow-up time. Categorical comparators were drawn as follows. Glioma: WHO 2021 primary classes.^30^ UCEC: TCGA molecular subtypes from Kandoth et al.^31^ with the PanCancer Atlas reannotation.^9^ BRCA: OncotypeDX 21-gene Recurrence Score (RS), reconstructed from TCGA RNA-seq via the Paik 2004 formula^32^ as released in the DLRS resource (Howard et al.,^33^ file `tcga_brca_complete.csv`, column `odx_train`). PRAD: 22-gene Decipher metastasis classifier,^34,35^ taken from the curatedPCaData uniform processing of TCGA-PRAD (Laajala et al.,^36^ ExperimentHub resource EH8024, `decipher` score in the `scores` slot). For BRCA and PRAD, the continuous score was tertile-binned within the analytic cohort to define Low / Intermediate / High strata; published vendor thresholds (e.g. Decipher 0.45/0.6) do not transfer to the published reconstructed score and were not used.

#### Risk-score derivation

For BRCA and PRAD, a degree-1 polynomial Cox proportional-hazards model with interaction on the two UMAP coordinates was fit against TCGA disease-specific survival truncated at 60 months for BRCA (DSS, 47 events at 5 years, n = 911) and against TCGA progression-free interval truncated at 60 months for PRAD (PFI, 62 events at 5 years, n = 391). PFI was used for PRAD per the Liu 2018 PanCancer Clinical Data Resource recommendation,^29^ because TCGA-PRAD has only ∼5 disease-specific death events and DSS is therefore underpowered. The polynomial-Cox fit yielded a continuous per-sample risk score (predicted partial hazard); three risk groups were defined by partitioning the continuous score with log-spaced thresholds across the score range (low, intermediate, high). Survival stratification was visualised with Kaplan–Meier curves, with a log-rank test across the three groups. Cross-tabulation of the three risk groups against the published transcriptomic comparator (OncotypeDX RS tertiles for BRCA, Decipher score tertiles for PRAD) documented the correspondence between the joint-genomic and transcriptomic stratifications.

#### Independence of the TESSERA score from the published transcriptomic comparator

For BRCA and PRAD we additionally fit a joint Cox proportional-hazards model with both predictors as standardised continuous covariates: hazard ∼ *z*(comparator) + *z*(log Risk_Score), where the comparator is the continuous OncotypeDX RS or Decipher score and Risk_Score is the TESSERA polynomial-Cox partial hazard. Both predictors were standardised to unit variance within the analytic cohort. Hazard ratios are reported per +1 SD with 95% Wald confidence intervals. To visualise stratification within the comparator’s most discretionary tier, we restricted to patients in the OncotypeDX-Intermediate (BRCA) or Decipher-Intermediate (PRAD) stratum and split them at the median TESSERA log Risk_Score, then plotted Kaplan–Meier curves and reported the within-stratum log-rank P value. The glioma and UCEC cohorts were analysed only by unsupervised projection and subtype-overlay visualisation and did not enter the polynomial-Cox risk-score analysis or the joint Cox.

### Counterfactual treatment-effect pipeline (DR-learner)

For each cohort (CRC FOLFOX vs FOLFIRI; PDAC FOLFIRINOX vs gemcitabine plus nab-paclitaxel) we estimated the per-patient treatment effect with a doubly-robust learner (DR-learner) on the TESSERA representation, a meta-learner formulation in which the second-stage regression target is constructed from the augmented inverse-probability-weighted (AIPW) pseudo-outcome and inherits the doubly-robust consistency property.^41,42^ Per-patient joint SNV+CNA features were obtained by passing each patient’s somatic variants and MSK-CHORD copy-number segments (raw, untrimmed; quantile-normalised onto the TCGA segment-mean distribution as above) through the TCGA-pretrained TESSERA model without retraining. Token-level SNV and segment-level CNA embeddings were aggregated by concatenating per-modality mean and max pools, log1p-transformed SNV and CNA counts were appended, and the resulting matrix was passed through a feature-engineering chain of `StandardScaler` → PCA (retaining 99% of variance) → `StandardScaler`, fit once per cohort.

#### Doubly-robust pseudo-outcome

For each patient the pseudo-outcome

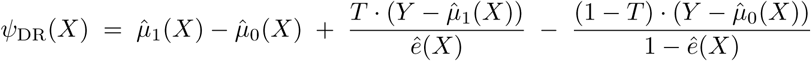

was constructed, where 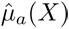 is the predicted restricted-mean PFS over a 36-month horizon under arm *a* from a Cox-PLS regression fit only on patients receiving arm *a*, 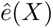 is the propensity score from a logistic-PLS regression fit on the full cohort, *T* is the binary treatment indicator (1 = arm 1, 0 = arm 0), and *Y* is the observed PFS time clipped at the 36-month horizon. Cox-RMST predictions were obtained by trapezoid-integrating the predicted Cox survival function on a Δ*t* = 0.5-month grid. Each nuisance model 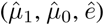 was fit by sparse partial-least-squares (SparsePLS) on the PCA-reduced features with separate hyperparameters per nuisance, and trained out-of-fold within each outer fold to remove training-set bias from the constructed pseudo-outcome. Propensity scores were clipped to [0.05, 0.95] before the AIPW combination to bound the inverse-probability weights.

#### Sparse-PLS regression on the pseudo-outcome

Within each outer fold, a sparse-PLS regression 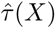 was fit on the held-out *ψ*_DR_ (the DR-learner second-stage regression) and used to score the held-out outer-test patients, producing fully out-of-fold per-patient 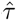 scores in months of Δ-PFS. The sign of sparse-PLS components is not determined by the optimisation, so the resulting 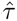 score has no built-in orientation. We sign-aligned 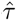 so that larger values correspond to larger predicted advantage of arm 1 by inverting the sign whenever the raw treatment-by-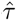 interaction Cox HR exceeded 1.

#### Outer/inner cross-validation

The full pipeline (feature-engineering chain, three nuisance regressions, propensity, sparse-PLS regression on *ψ*_DR_) was wrapped in a 10-fold outer / 5-fold inner stratified nested cross-validation. The outer fold was used only for held-out scoring; the inner fold was used to construct the per-patient nuisance predictions. Outer folds were stratified on the joint (arm, event) label so each fold was balanced on both treatment and outcome; inner folds were stratified on event alone, since each arm-restricted nuisance holds arm constant within its split, and the propensity inner fold balances outcome rather than the variable being modelled. The horizon for RMST was 36 months throughout, matching the administrative censoring horizon used for the survival visualisations.

#### Indifference threshold

The per-cohort indifference threshold 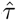_0_ was defined from the treatment-by-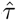 interaction Cox model fit on PFS: log HR(arm | 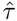) = *β_T_* + *β*_int_ · 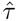, solved for unit hazard, 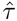_0_ = −*β_T_ /β*_int_. Patients with 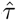 *>* 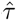_0_ were classed predicted-arm-1-favoured; those with 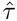 *<* 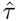_0_ predicted-arm-0-favoured. The same interaction Cox was used to evaluate the smooth analytic HR(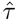) curve on a fine grid for visualisation.

#### Stratified survival analyses

For each cohort and each endpoint (PFS as the training endpoint; OS as a held-out validation endpoint), the within-stratum cross-arm hazard ratio was fit by Cox time ∼ arm on each predicted-favoured subset. Pooled-cohort splits tested the prognostic-vs-predictive identity by Cox PFS ∼ **1**[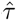 *>* 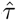_0_] on the full cohort regardless of arm; a HR close to 1 there is the prognostic-null prerequisite for interpreting the per-stratum cross-arm effects as predictive rather than prognostic. Kaplan–Meier curves used `lifelines` step estimates with administrative censoring at the cohort horizon (36 months for PFS, 60 months for OS).

#### Feature-slice ablation

To localise where in the foundation-model representation the predictive signal lives, the same DR-learner pipeline was re-run on three nested feature subsets of the per-patient embedding: a Local subset using only the local-attention SNV and CNA features (variant- and segment-intrinsic features); a Global subset additionally including the within-modality global-attention features (variant-variant and segment-segment interactions); and a Full subset additionally including the InfoNCE-aligned cross-modal features (the representation used in the main analysis). Each subset is feature-engineered, nuisance-fit, and DR-second-stage-fit independently, yielding its own out-of-fold 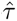 and 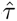_0_. The per-stratum cross-arm Cox hazard ratios under each feature subset were tabulated for both cohorts to identify which representational components contribute the predictive signal.

### Genomic signature decomposition

To localise the genomic basis of the CRC predictive direction, we composed the post-aggregation chain of the DR-learner into a single effective linear map and decomposed the resulting per-(patient, feature) attribution matrix into a sparse low-rank signature structure.

#### Effective coefficients

The DR-learner second-stage regression 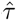(*X*) is linear in *X* once the chain `StandardScaler` → `PCA`(0.99) → `StandardScaler` → `SparsePLS` is composed, so all post-aggregation transforms collapse to a single 3,716-dimensional effective coefficient vector *β*_eff_ such that 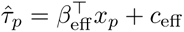 for every patient *p*. The pipeline was re-fit once on the full cohort with no outer split (held-out predictive performance is already validated under the nested cross-validation described above) and the composition was verified by a per-patient reconstruction check, requiring 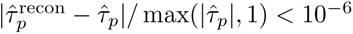 across all patients in both cohorts.

#### Per-(patient, variant) attributions

Mean-pool features are linear in their inputs: variant *v*’s share of 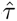*_p_* from the SNV mean pool is 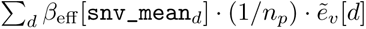, where 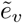 is the RobustScaled SNV embedding of variant *v* and *n_p_* is patient *p*’s variant count. Max-pool features are not linear, but each max-pool dimension is a per-patient argmax over variants, so variant *v* contributes only at dimensions *d* where *v* is the argmax: 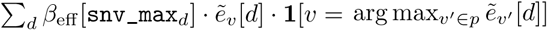. The per-(patient, variant) SNV attribution is the sum of the two contributions, signed in months of Δ-PFS. The same logic applied to CNA segments via the cna_mean and cna_max coefficients. The two patient-level burden columns (log(1+TMB) and log(1+nCNA)) yielded a single per-patient scalar contribution. The reconstruction Σ*_v_* attribution*_pv_* + burden*_p_* + *c*_eff_ = 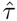*_p_* was verified to |Δ|*/* max(|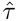|, 1) *<* 10^−4^ for every patient.

#### Feature-level aggregation

SNV attributions were aggregated to gene symbols via the `Hugo_Symbol` field. CNA segment attributions were aggregated to chromosomal arms (p and q for chromosomes 1–22 plus X and Y, 42 arms) by interval-overlap: for each (segment, arm) pair with non-zero overlap in base pairs, the segment’s attribution was credited to the arm. Arm-level rather than gene-level resolution is the appropriate target for panel CNA because the median MSK-IMPACT segment spans roughly 29 Mb, well above gene scale. The result is a per-(patient, feature) attribution matrix *A* ∈ ℝ*^P^* ^×^*^G^* with *G* = 420 features for CRC (378 SNV genes plus 42 CNA arms) and *G* = 420 features for PDAC.

#### Sparse signature decomposition

We decomposed *A* via the penalised matrix decomposition of Witten et al.:^46^ *A* ≈ *UDV* ^⊤^ with ‖*U*_·_*_k_*‖_2_ ≤ 1, ‖*V*_·_*_k_*‖_2_ ≤ 1, ‖*U*_·_*_k_*‖_1_ ≤ *c_u_*, and ‖*V*_·_*_k_*‖_1_ ≤ *c_v_*. The patient-side budget was set to 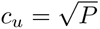 (no penalty on patient loadings); the feature-side budget was set to 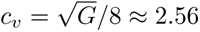 (moderate sparsity). The rank was fixed at *K* = 5 for the CRC analysis. PMD components are signed but their global sign is not identified by the optimisation; we did not enforce a global sign convention on *V* at the decomposition step, but downstream in the heatmap visualisation each cell value was rendered as *v_g,k_* for arm features (the row label encodes sign(*ρ_g_*) as AMP or LOSS) and as *v_g,k_* · sign(*ρ_g_*) for SNV features (where the row label is fixed to MUT and the directional sign is folded into the cell value), so that positive cells uniformly denote features whose labelled biological direction (AMP, LOSS, or MUT) contributes positively to the FOLFOX axis.

#### Arm-direction map

Each CNA-arm column of *A* was annotated with a biology direction (amplification or loss) by computing the Spearman correlation *ρ_g_* between the column and the per-patient length-weighted mean Segment_Mean over segments overlapping that arm. Positive *ρ_g_* marks an amplification-driven attribution direction (high attribution corresponds to elevated copy number on that arm); negative *ρ_g_*marks a loss-driven direction. The same Spearman procedure was applied to SNV columns against the per-patient mutation indicator for each gene, yielding the SNV *ρ_g_*used in the heatmap sign convention above.

#### TP53/KRAS/17p simple-rule subgroups

TP53 and KRAS were binarised at any coding-impact mutation. Per-patient 17p loss was defined as the length-weighted mean Segment_Mean across CNA segments overlapping the 17p arm (hg19 boundaries 0–22.2 Mb), thresholded at log_2_ *<* −0.2. Five exclusive subgroups were enumerated: TP53^+^/KRAS^+^/17p^−^ (both-mutated with 17p loss), TP53^+^/KRAS^+^ (both-mutated, regardless of 17p), TP53^+^/KRAS^−^ (TP53-only), TP53^−^/KRAS^+^ (KRAS-only), and TP53^−^/KRAS^−^ (double-wildtype). Per-subgroup arm hazard ratios were fit by Cox PFS ∼ arm within each mask, with degenerate strata (*n <* 10, single-arm, or fewer than 3 events) reported as not estimable.

#### PDAC simple-rule subgroup

The PDAC triplet was derived from Sig 8 of the rank-10 PMD decomposition. The top three signed loadings in Sig 8 are 10q amplification, TP53 coding mutation, and 17p loss; 10q-amp prevalence in the analytic PDAC cohort is ∼2.5% (*n* = 19), too low to support a stable subgroup, so 20q amplification (∼21% prevalence) was substituted as the closest concordant amplification axis on the same signature. The published triplet (TP53 mutated, 17p intact at log_2_ ≥ −0.2, 20q amplified at log_2_ *>* +0.2) reflects this substitution.

### DepMap cell-line validation

Cancer cell-line somatic mutations and copy-number segments (with LoH annotations) were obtained from the Cancer Cell Line Encyclopedia / DepMap 24Q2 release.^47^ Segment coordinates were lifted hg38→hg19 with `pyliftover`; SNVs were filtered to single-base substitutions and converted to TCGA MAF format. To match the assay coverage on which 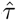 was fitted, both modalities were restricted to MSK-IMPACT505 panel coverage (SNVs to the 505 panel genes; CNA segments to those overlapping any panel gene region) and subsampled per cell line to at most 1,000 variants and 1,000 segments using the same priority logic as TCGA pretraining. DepMap segment-mean values were quantile-normalised onto the TCGA training distribution before inference, matching the MSK-CHORD CNA normalisation procedure described above. Frozen TESSERA produced per-token embeddings, the MSK-CHORD-fitted SNV and CNA RobustScalers were applied, mean and max pools were computed per cell line, and 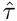 was evaluated as *β*_eff_ · *x* + *c*_eff_ using the saved coefficients from the MSK-CHORD CRC fit.

CTRPv2^48^ dose-response area-under-the-curve values were used for oxaliplatin (the FOLFOX-specific cytotoxic) and SN-38 (the active metabolite of irinotecan, the FOLFIRI-specific cytotoxic); 5-FU and leucovorin are shared by both regimens and therefore cannot drive a FOLFOX-vs-FOLFIRI selection signal. For colorectal cell lines with both drugs measured (*n* = 44), AUCs were z-scored within the cohort independently per drug and converted to sensitivities by negation (lower CTRPv2 AUC indicates greater sensitivity). The FOLFOX-over-FOLFIRI preference was computed as the difference of the two z-scored sensitivities (positive values ⇒ greater oxaliplatin than SN-38 sensitivity). Spearman’s *ρ* between 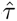 and the preference was reported with a one-sided P-value reflecting the predicted positive direction (high 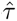 ⇒ FOLFOX-favoured prediction ⇒ greater oxaliplatin sensitivity).

### FOLFOXai comparator replication

The FOLFOXai signature^56^ was re-implemented on the MSK-CHORD first-line stage IV CRC FOLFOX-vs-FOLFIRI cohort used for the DR-learner pipeline above. Because FOLFOXai’s labelling endpoint (TTNTD) differs from the DR-learner’s (PFS), the FOLFOXai evaluation cohort comprises 1,443 TTNTD-evaluable patients versus 1,452 PFS-evaluable patients in the DR-learner; the 9-patient gap reflects PFS-evaluable patients without a recorded next-line switch. Features were the Abraham 67-gene panel: for each gene, a `_mutations` indicator (1 if the patient carries any coding-impact SNV at that gene, 0 otherwise) and a `_copies` value (length-weighted mean of 2 · 2^Segment_Mean^ clipped at ±3, default diploid 2.0 when missing). Of the 67 panel genes, 28 are profiled by MSK-IMPACT and contribute non-zero features; the remaining 39 are absent from MSK-CHORD calls and enter as zero (no SNV) and diploid (Segment_Mean unobserved). Patients were labelled IB if their time to next treatment discontinuation (TTNTD) was ≥ 270 days and DB if TTNTD was *<* 270 days with the event observed, matching Abraham et al.; FOLFOX patients with TTNTD *<* 270 days and the event censored were excluded from training as short-follow-up. The classifier was a 5-configuration ensemble of 1,000-tree balanced-class `RandomForestClassifier`s (`max_features` in {`sqrt`, `log2`, 0.3, 0.5}, `min_samples_leaf` in {5, 10, 20}, random states 0–4). The ensemble-mean *p*_IB_ was computed per patient; the 3% no-call buffer (0.47 ≤ *p*_IB_ ≤ 0.53) was applied per the original paper. FOLFOX out-of-fold predictions used 5-fold stratified cross-validation (seed 42); FOLFIRI patients were scored out-of-sample by fitting each configuration on the full FOLFOX IB/DB cohort and applying it to the FOLFIRI patients. Although FOLFOXai is trained on a binary TTNTD threshold within the FOLFOX arm, downstream evaluation uses continuous PFS and OS on the full cohort, matching both the endpoint set used for TESSERA’s DR-learner above (so the two methods are compared on the same outcomes) and the PFS and OS endpoints reported for clinical validation in the original Abraham analyses. Three downstream survival analyses were computed under the same TESSERA-canonical endpoint construction described above (36-month administrative censor for PFS, 60-month for OS): a pooled IB-versus-DB Kaplan–Meier on the full no-call-excluded cohort, and arm-stratified FOLFOX-versus-FOLFIRI Kaplan–Meier separately within the predicted-IB and predicted-DB strata. Per-analysis statistics were the Cox proportional-hazards hazard ratio with 95% Wald CI and the two-sided logrank *P*.

### Statistical analysis and software

All performance metrics were computed with 95% confidence intervals using bootstrap resampling (n = 1,000 iterations). Survival models and tests used the `lifelines` (v0.29) Python package. Pretraining was performed in Python 3.11 with TensorFlow 2.20 on NVIDIA RTX 6000 Ada GPUs; downstream survival and treatment-effect analyses used scikit-learn, pandas, numpy, and scipy. Visualisations were generated with matplotlib and seaborn using colourblind-safe palettes.

### Data availability

#### Pretraining and TCGA-derived analyses

The TCGA Pan-Cancer Atlas somatic mutation (MC3) MAF and ABSO-LUTE copy-number segment files are publicly available from the GDC Pan-Cancer Atlas publication page (https://gdc.cancer.gov/about-data/publications/pancanatlas). The TCGA Pan-Cancer Clinical Data Resource is provided as the supplementary release accompanying Liu et al.^29^ (`TCGA-CDR-SupplementalTableS1.xlsx`). The GRCh37 reference genome (assembly `GCF_000001405.25_GRCh37.p13`) was obtained from NCBI (https://ftp.ncbi.nlm.nih.gov/genomes/all/GCF/000/001/405/GCF_000001405.25_GRCh37.p13/).

#### Variant pathogenicity prediction

ClinVar variant annotations are publicly available from NCBI (https://www.ncbi.nlm.nih.gov/clinvar/); the GRCh37 VCF release used in this study is available from https://ftp.ncbi.nlm.nih.gov/pub/clinvar/vcf_GRCh37/.

#### Cross-platform validation

The AACR Project GENIE v18.0 public release is available from Synapse (accession [*syn7222066*], https://www.synapse.org/Synapse:syn7222066). The MSK-CHORD 2024 release is available through cBioPortal (study `msk_chord_2024`, https://www.cbioportal.org/study/summary?id=msk_chord_ `2024`) and Synapse as described in Jee et al..^28^

#### Prognostic-cohort molecular-subtype comparators

The WHO 2021 glioma reclassification table is provided as supplementary material to Leiria et al.^30^ and is also available at https://github.com/sysbiomed/MONET/blob/master/Final-outputs/Matrix_WHO2021.csv. TCGA UCEC PanCancer Atlas molecular subtype labels (Kandoth et al.^31^ with the Hoadley et al. reannotation^9^) are available from cBioPortal (study `ucec_tcga_pan_can_atlas_2018`, https://www.cbioportal.org/study/clinicalData?id=ucec_tcga_pan_can_atlas_2018). DLRS-reconstructed OncotypeDX RS and MammaPrint scores for TCGA-BRCA are released by Howard et al.^33^ at https://github.com/fmhoward/DLRS. curatedPCaData transcriptomic scores (Decipher, OncotypeDX GPS, Prolaris, AR-signaling) for TCGA-PRAD are available as Bioconductor ExperimentHub resource `EH8024`^36^ (https://bioconductor.org/packages/curatedPCaData/).

#### Cell-line drug-response validation

Cancer Cell Line Encyclopedia / DepMap 24Q2 somatic mutations, copy-number segments, and clinical metadata^47^ are available from https://depmap.org/portal/download/all/. Cancer Therapeutics Response Portal v2 (CTRPv2) drug-response area-under-the-curve values for oxaliplatin and SN-38^48^ are available from the Broad Institute portal (https://portals.broadinstitute.org/ctrp.v2.2/).

### Code availability

#### Source code

All code for model pretraining, downstream analyses, and figure generation is available at https://github.com/JW-Sidhom-Lab/tessera under the PolyForm Noncommercial Licence 1.0.0. The TESSERA foundation-model package is also published on the Python Package Index as `tessera-foundation` (`pip install tessera-foundation`); the package wraps both the inference API used in this study and the training entry points, and downloads the pretrained weights on demand.

#### Pretrained foundation-model weights

Pretrained weights for both joint SNV+CNA variants (with and without the loss-of-heterozygosity head) are released on the Hugging Face Hub at https://huggingface.co/JW-Sidhom-Lab/tessera-foundation under a Creative Commons Attribution-NonCommercial 4.0 International (CC-BY-NC-4.0) licence and are downloaded automatically by tessera.featurize() on first use.

#### TCGA Pan-Cancer Atlas TESSERA representations

Per-sample and per-token TESSERA SNV and CNA features for the full TCGA Pan-Cancer Atlas cohort (∼10,000 tumours), computed under the canonical joint SNV+CNA InfoNCE-aligned model used throughout this study, are deposited as a permanent HDF5 archive at Zenodo (DOI: `10.5281/zenodo.20419467`). The deposit provides a directly reusable representation of the TCGA Pan-Cancer Atlas, so that downstream foundation-model-based analyses can build on TESSERA without rerunning pretraining or inference. Format, schema, and citation details are documented in the deposit README.

## Acknowledgements

This work was supported by the Weill Cornell × Mastercard AI / Big Data Seed Grant. The results here are in whole or part based on data generated by the TCGA Research Network, the AACR Project GENIE consortium, and the MSK-CHORD project.

During manuscript preparation, the authors used Claude Code (Anthropic) to refine the language of the manuscript and to review the analysis code for clarity, correctness, and reproducibility, and Claude Design (Anthropic) to produce the schematic architecture figures presented in the manuscript. All scientific content, analyses, final language choices, and final figure compositions remain the authors’ responsibility; every suggested change was reviewed and verified by the authors before adoption.

## Author contributions

J.-W.S. conceived the study, developed the foundation-model architecture and pretraining procedures, performed all analyses, prepared the figures, and wrote the manuscript. A.S.B. contributed to the development of the methods and to the choice of downstream applications. O.E. advised on the computational methods and ensured the rigour of the analyses. M.A.S. shaped the clinical scope and motivation of the work, and was closely involved in the design and interpretation of the translational analyses. A.S.B., O.E., and M.A.S. supervised the project and revised the manuscript. All authors read and approved the final manuscript.

## Additional information

**Supplementary Information** is available for this paper.

**Correspondence and requests for materials** should be addressed to J.-W.S.

