## Supplementary Data for "A Foundation Model for the Cancer Genome"

### A Foundation Model for the Cancer Genome - Supplementary Information

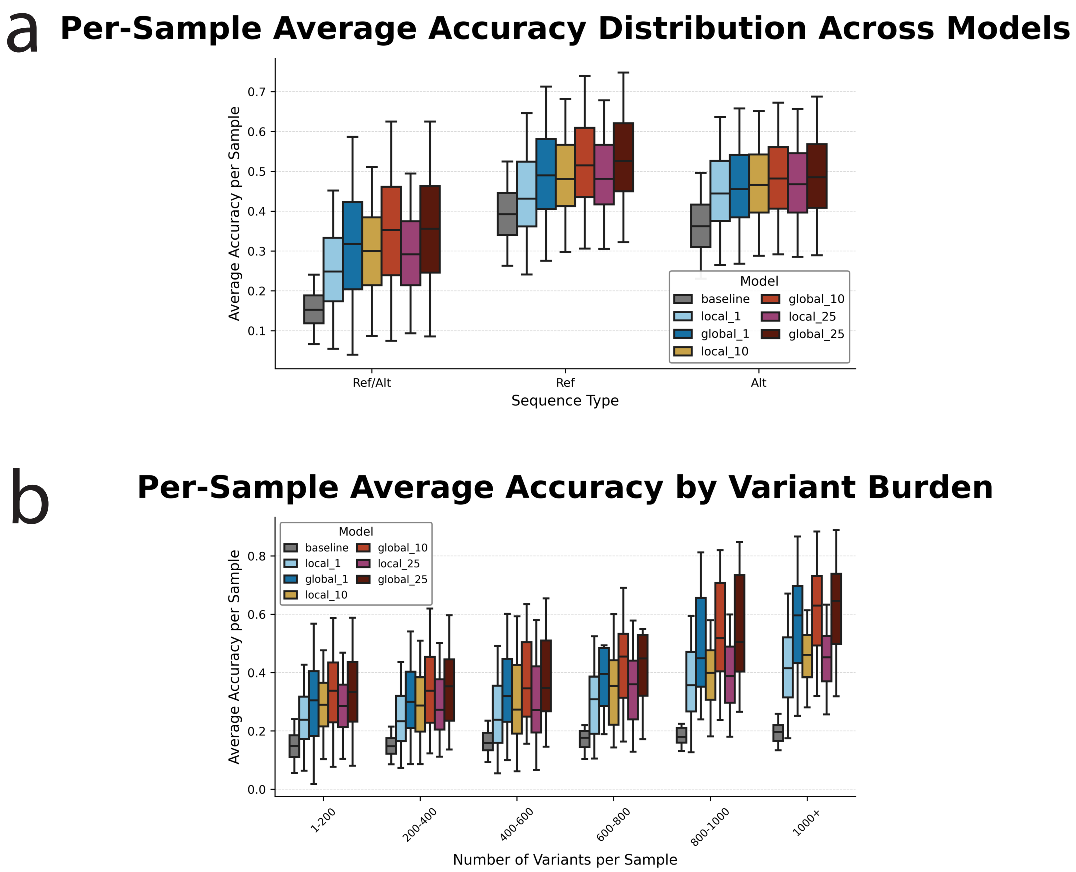


**Supplementary Figure 1. Sample-level accuracy distributions for masked token prediction.** Masked token prediction accuracy aggregated at the sample level (mean accuracy across all variants in each sample) on TCGA WES validation data. **a,** Distribution of per-sample average accuracy across model architectures, stratified by prediction type (Ref/Alt, Ref only, Alt only). Box plots show median, quartiles, and outliers for each model variant: baseline, local attention with 1bp, 10bp, 25bp context, and global attention with 1bp, 10bp, 25bp context. **b,** Distribution of per-sample average accuracy stratified by variant burden (number of variants per sample). Each box represents the distribution of sample-level accuracies within each variant burden bin. Variant burden bins: 1-200, 200-400, 400-600, 600-800, 800-1000, >1000 variants per sample.

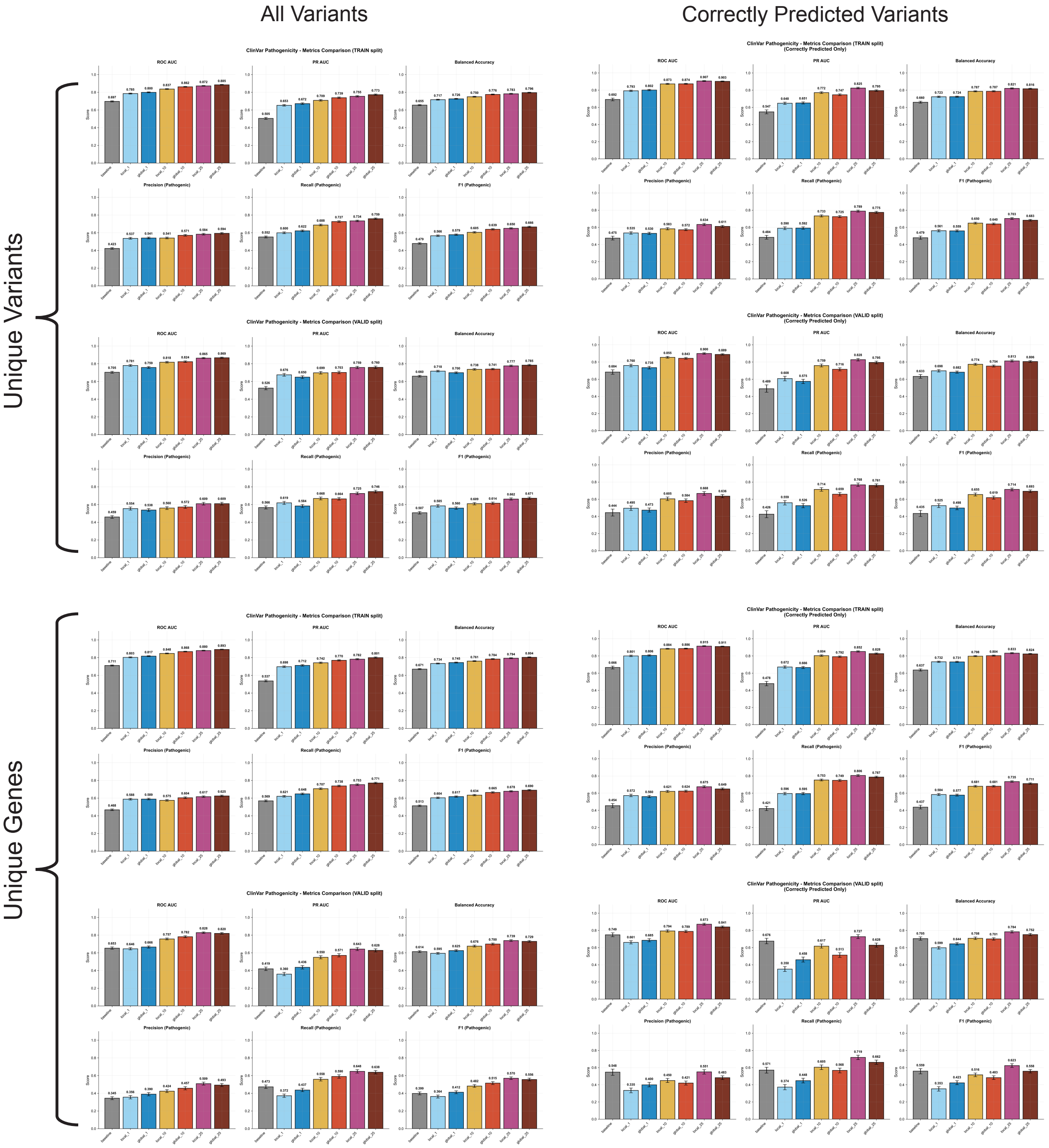


**Supplementary Figure 2. Comprehensive variant effect prediction performance metrics.** Detailed evaluation of ClinVar pathogenicity prediction models across multiple metrics, evaluation strategies, and data subsets. The figure is organised as a 2 × 2 grid with columns representing data subsets (all variants vs. correctly predicted variants) and rows representing evaluation strategies (unique variants vs. unique genes). “Correctly predicted variants” refers to the subset of variants where the model correctly predicted masked reference and alternate alleles during self-supervised pretraining, testing whether variants the model “understands” at the sequence level yield more reliable pathogenicity predictions. **Top row (Unique Variants):** Standard variant-level deduplication where test variants (unique chr:pos:ref:alt combinations) are completely held out from training. Left panel shows performance metrics on all held-out test variants. Right panel shows metrics restricted to correctly predicted variants. **Bottom row (Unique Genes):** Gene-level deduplication where test variants come from genes completely absent during training, testing generalisation to novel genomic contexts. Left panel shows metrics on all held-out variants from unseen genes. Right panel shows metrics on the subset of correctly predicted variants from unseen genes. Within each panel, six performance metrics are shown: ROC AUC, PR AUC, balanced accuracy, precision, recall, and F1 score. Each metric compares model variants: baseline (no sequence context), local attention at 1bp, 10bp, and 25bp, and global attention at 1bp, 10bp, and 25bp context. Bar plots show mean metric values with error bars. Each quadrant is further subdivided vertically into a TRAIN-split sub-panel (top, training-set performance) and a VALID-split sub-panel (bottom, held-out validation performance). The VALID-split metrics correspond to the values reported in the main text and in Figure 1h–o.

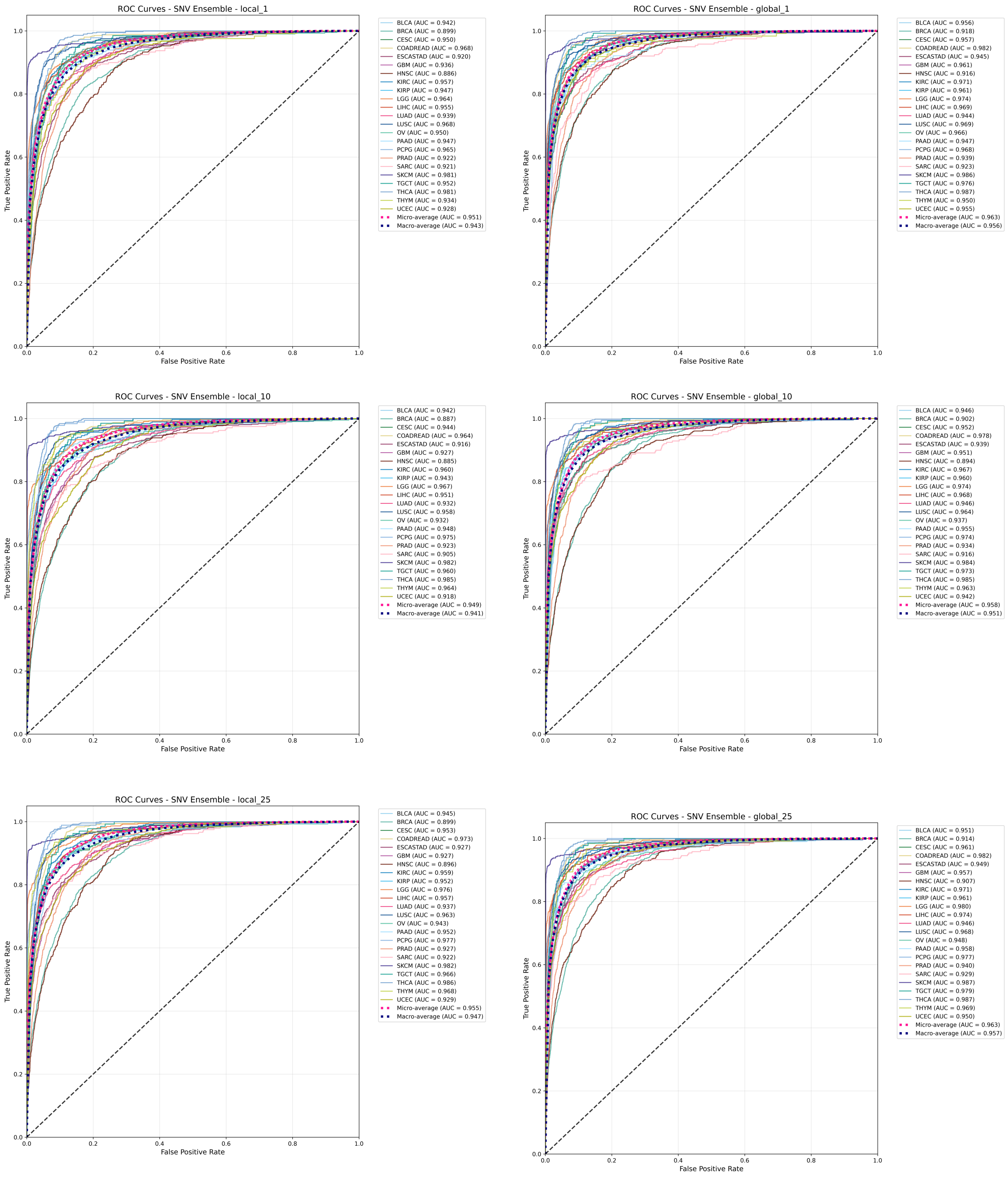


**Supplementary Figure 3. Per-class ROC curves for SNV-based tumour type classification across model architectures and context sizes.** Receiver operating characteristic (ROC) curves showing SNV model classification performance for each tumour type (class) across all model configurations. The figure is organised as a $3 \times 2$ grid where columns represent attention mechanisms (local vs. global) and rows represent sequence context sizes (1bp, 10bp, 25bp). **Column 1 (Local Models):** ROC curves for SNV models using local attention mechanisms at 1bp (top), 10bp (middle), and 25bp (bottom) context windows. **Column 2 (Global Models):** ROC curves for SNV models using global attention mechanisms at 1bp (top), 10bp (middle), and 25bp (bottom) context windows. Each panel displays individual ROC curves for all tumour type classes, allowing comparison of per-class SNV-based classification performance across different model configurations.

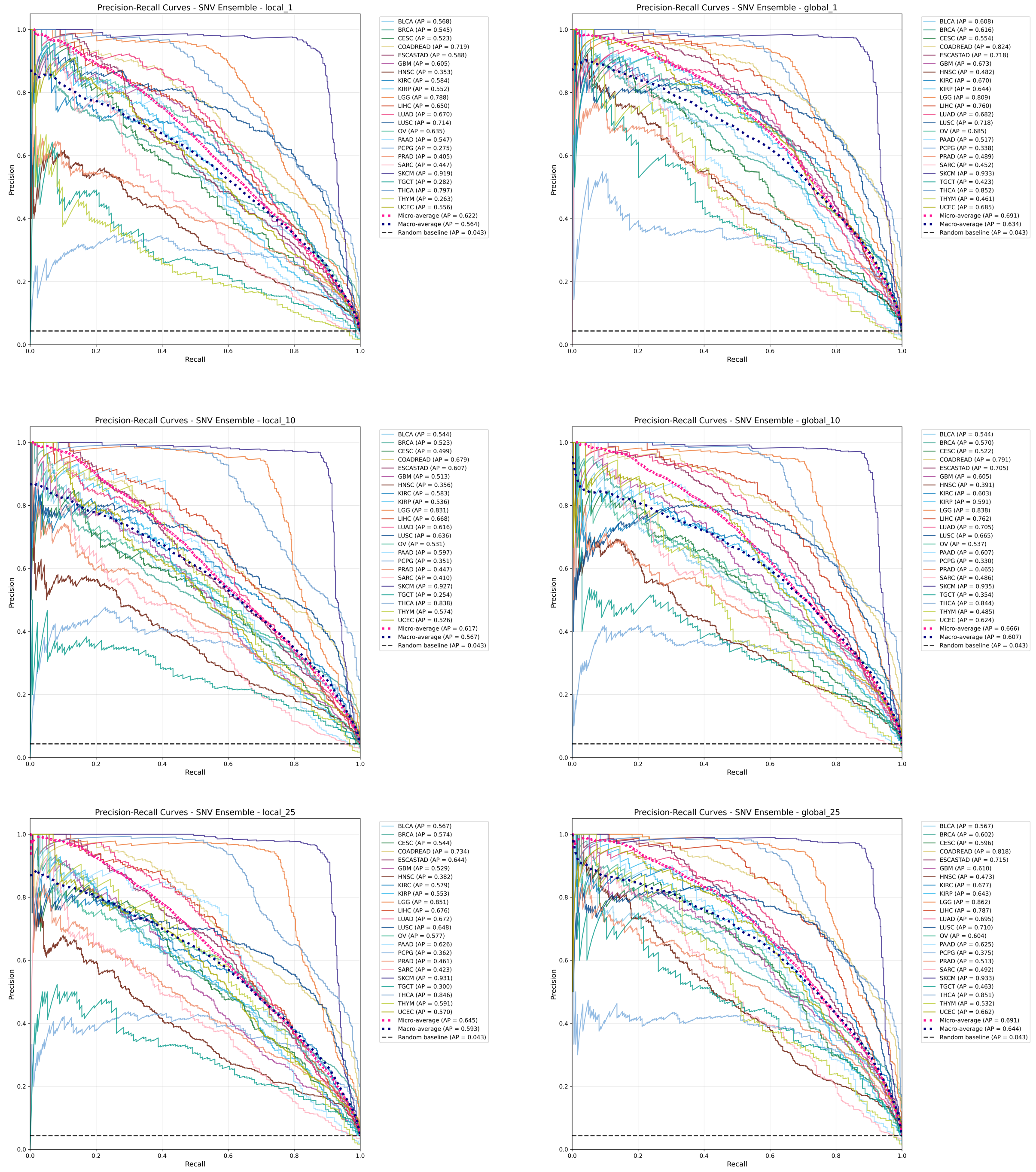


**Supplementary Figure 4. Per-class precision-recall curves for SNV-based tumour type classification across model architectures and context sizes.** Precision-recall (PR) curves showing SNV model classification performance for each tumour type (class) across all model configurations. The figure is organised as a $3 \times 2$ grid where columns represent attention mechanisms (local vs. global) and rows represent sequence context sizes (1bp, 10bp, 25bp). **Column 1 (Local Models):** PR curves for SNV models using local attention mechanisms at 1bp (top), 10bp (middle), and 25bp (bottom) context windows. **Column 2 (Global Models):** PR curves for SNV models using global attention mechanisms at 1bp (top), 10bp (middle), and 25bp (bottom) context windows. Each panel displays individual PR curves for all tumour type classes, allowing comparison of per-class SNV-based classification performance across different model configurations.

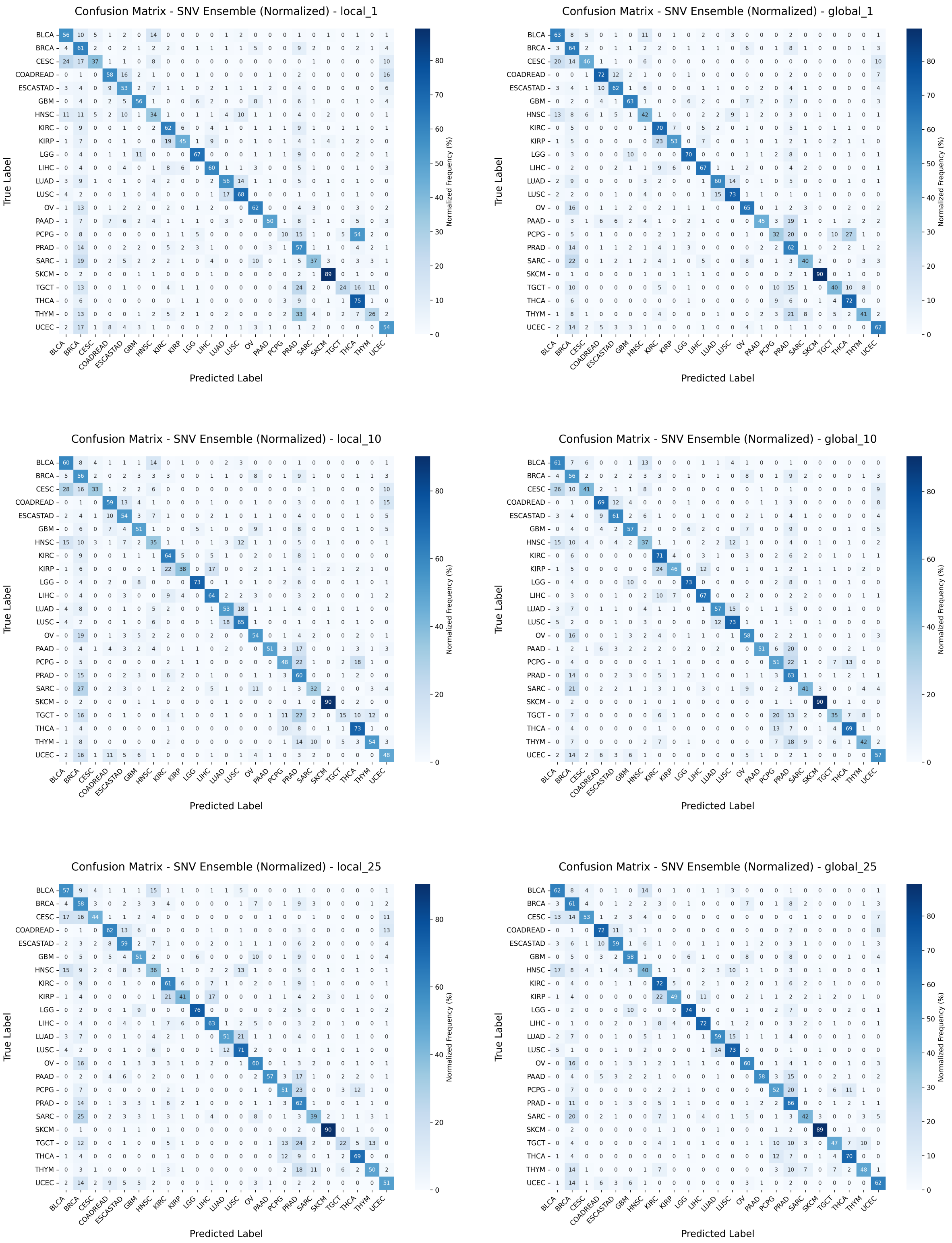


**Supplementary Figure 5. Normalised confusion matrices for SNV-based tumour type classification across model architectures and context sizes.** Normalised confusion matrices showing SNV model classification performance for each tumour type across all model configurations. The figure is organised as a $3 \times 2$ grid where columns represent attention mechanisms (local vs. global) and rows represent sequence context sizes (1bp, 10bp, 25bp). **Column 1 (Local Models):** Confusion matrices for SNV models using local attention mechanisms at 1bp (top), 10bp (middle), and 25bp (bottom) context windows. **Column 2 (Global Models):** Confusion matrices for SNV models using global attention mechanisms at 1bp (top), 10bp (middle), and 25bp (bottom) context windows. Each panel displays a normalised confusion matrix with true labels on the vertical axis and predicted labels on the horizontal axis, with values representing the proportion of samples from each true class assigned to each predicted class.

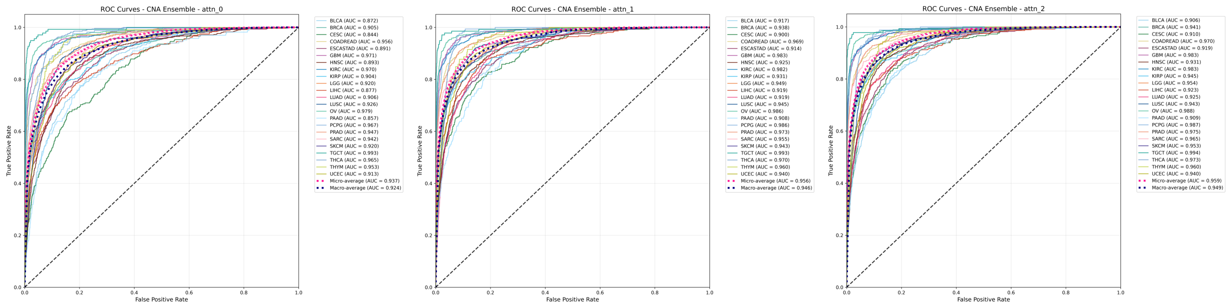


**Supplementary Figure 6. Per-class ROC curves for CNA-based tumour type classification across self-attention configurations.** Receiver operating characteristic (ROC) curves showing CNA model classification performance for each tumour type across all model configurations (attn_0, attn_1, attn_2). Each panel displays individual ROC curves for all tumour type classes.

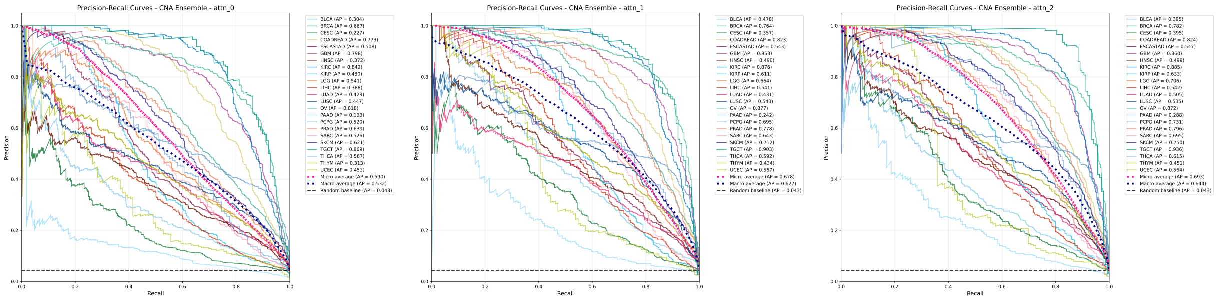


**Supplementary Figure 7. Per-class precision-recall curves for CNA-based tumour type classification across self-attention configurations.** Precision-recall (PR) curves showing CNA model classification performance for each tumour type across all model configurations (attn_0, attn_1, attn_2). Each panel displays individual PR curves for all tumour type classes.

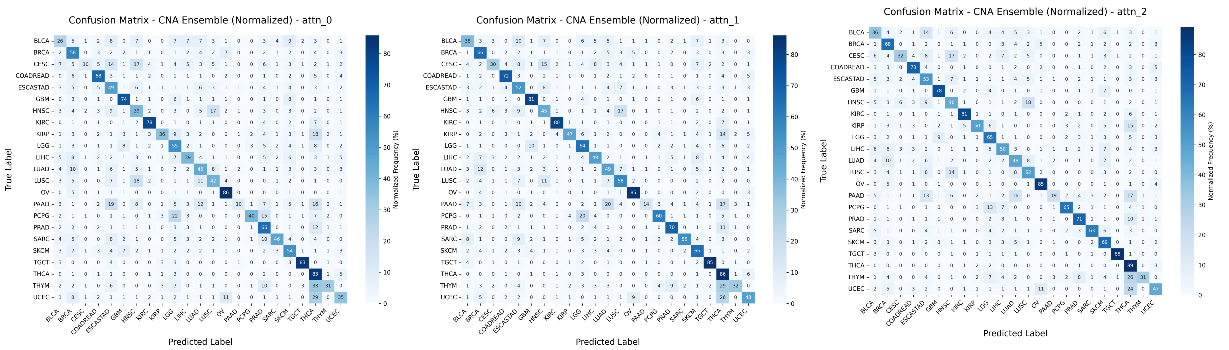


**Supplementary Figure 8. Normalised confusion matrices for CNA-based tumour type classification across self-attention configurations.** Normalised confusion matrices showing CNA model classification performance for each tumour type across all model configurations (attn_0, attn_1, attn_2). Each panel displays a normalised confusion matrix with true labels on the vertical axis and predicted labels on the horizontal axis, with values representing the proportion of samples from each true class assigned to each predicted class.

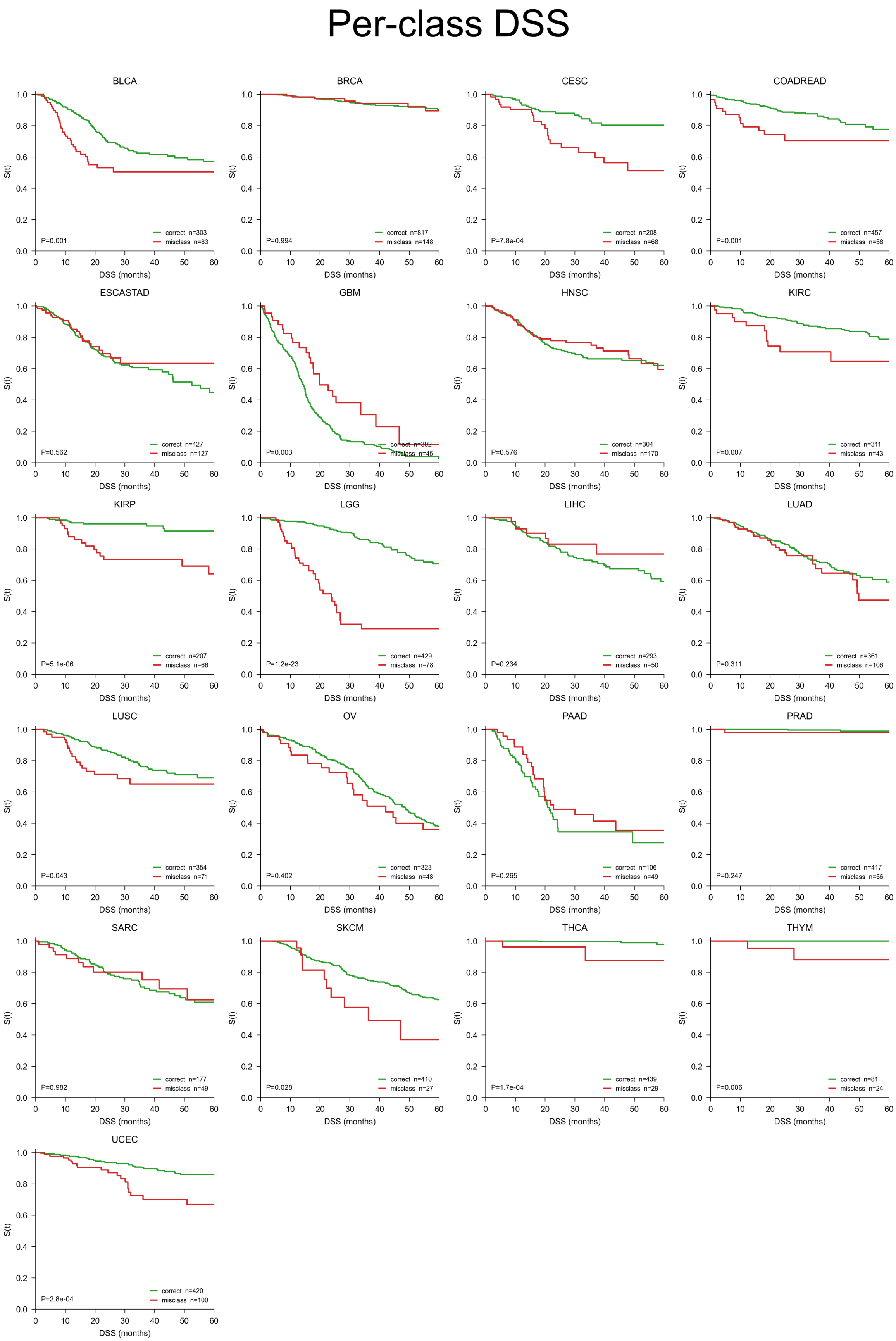


**Supplementary Figure 9. Per-class Kaplan–Meier curves of correctly classified versus misclassified TCGA tumours.** Small-multiples grid corresponding to the per-class forest plot in Fig. 4e of the main text. One panel per pathology class with at least 20 samples in each arm. Correctly classified samples shown in green; misclassified samples shown in red. Per-class logrank $P$-value annotated in each panel. Survival endpoint is TCGA disease-specific survival (DSS) with 60-month administrative censoring, matching the convention used throughout the predictive-biomarker analyses.

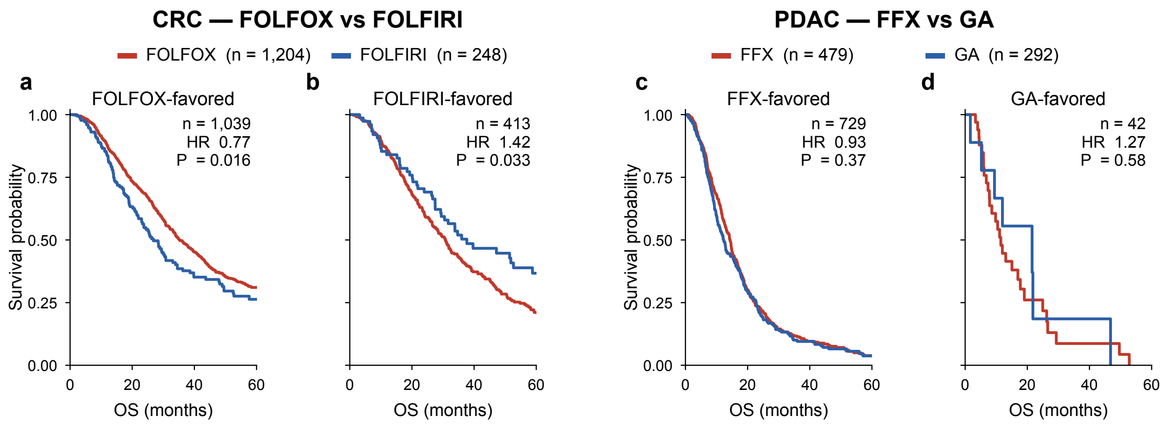


**Supplementary Figure 10. Overall-survival Kaplan–Meier curves within each predicted-favoured stratum defined by the DR-learner pipeline.** For each cohort, within-stratum cross-arm OS Kaplan–Meier curves are shown in the predicted-arm-1-favoured ($\hat{\tau} > \hat{\tau}_0$) and predicted-arm-0-favoured ($\hat{\tau} < \hat{\tau}_0$) subsets defined in Fig. 6. OS is truncated at the 60-month administrative censoring horizon. **a,** CRC predicted FOLFOX-favoured stratum: FOLFOX (red) versus FOLFIRI (blue). **b,** CRC predicted FOLFIRI-favoured stratum, colours as in (a). **c,** PDAC predicted FOLFIRINOX-favoured stratum: FOLFIRINOX (red) versus gemcitabine plus nab-paclitaxel (blue). **d,** PDAC predicted gemcitabine plus nab-paclitaxel-favoured stratum, colours as in (c). In each panel, $n$ denotes the within-stratum sample size, HR is the Cox proportional-hazards estimate of arm 1 versus arm 0 with 95% Wald CI, and $P$ is the associated Wald $P$-value.

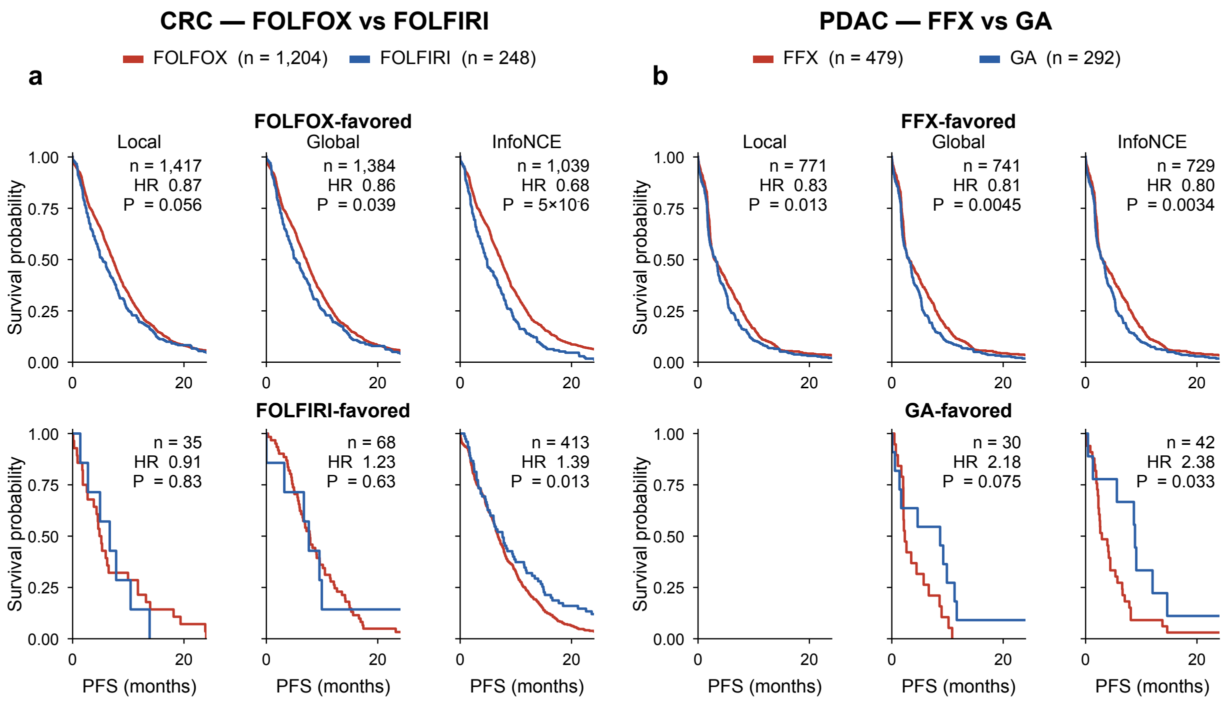


**Supplementary Figure 11. Feature-slice ablation of the DR-learner pipeline in CRC and PDAC.** The DR-learner pipeline of Fig. 6a was re-fit on three nested feature subsets of the per-patient embedding: *Local* (local-attention SNV and CNA features only), *Global* (additionally including within-modality global-attention features), and *InfoNCE* (additionally including the cross-modal InfoNCE-aligned features, the representation used in the main analysis). Each subset was fit, scored, and thresholded independently, yielding its own out-of-fold $\hat{\tau}$ and indifference threshold $\hat{\tau}_0$. PFS is displayed to 24 months for legibility, with administrative censoring at 36 months. **a,** CRC. Two rows of three PFS Kaplan–Meier panels: top row, predicted FOLFOX-favoured stratum; bottom row, predicted FOLFIRI-favoured stratum. Within each panel, FOLFOX is plotted in red and FOLFIRI in blue. Columns correspond to the Local, Global, and InfoNCE feature subsets. **b,** PDAC. Layout as in (a), with FOLFIRINOX (red) and gemcitabine plus nab-paclitaxel (blue). Top row, predicted FOLFIRINOX-favoured stratum; bottom row, predicted gemcitabine plus nab-paclitaxel-favoured stratum. In each panel, $n$ denotes the within-stratum sample size at that feature subset's $\hat{\tau}_0$, HR is the Cox proportional-hazards estimate of arm 1 versus arm 0, and $P$ is the associated Wald $P$-value.

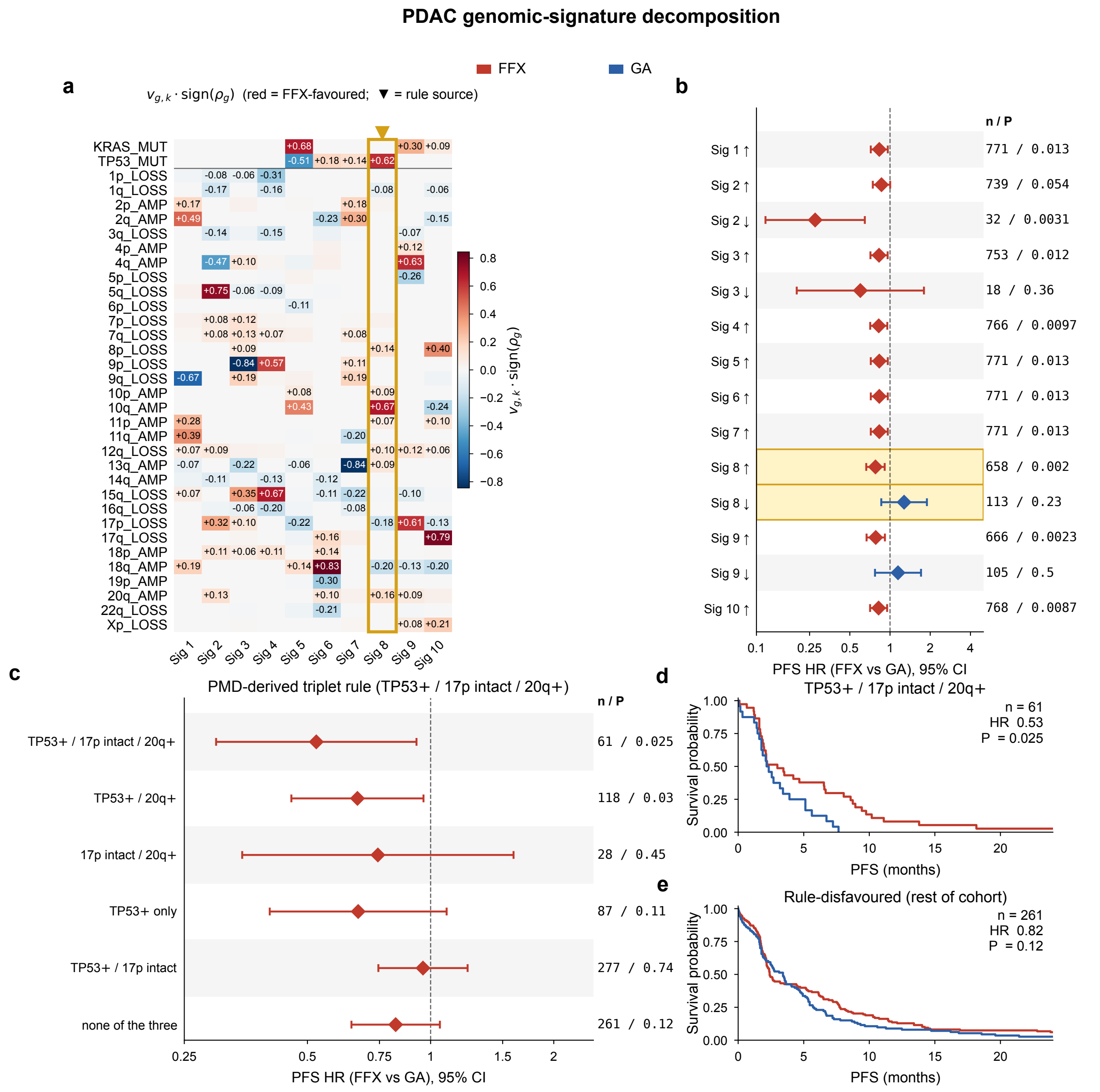


**Supplementary Figure 12. PDAC genomic-signature decomposition.** PMD (Fig. 6j–m) applied to the PDAC per-(patient, feature) attribution matrix at rank $K = 10$. Throughout: FFX = FOLFIRINOX, GA = gemcitabine plus nab-paclitaxel. **a,** $K = 10$ PMD signature heatmap. Rows: top-loaded features per signature; columns: Sig 1–10. Cell colour is the directional loading $v_{g,k} \cdot \mathrm{sign}(\rho_g)$ (red = FFX-favoured axis); row labels encode AMP or LOSS for arms, MUT for SNVs. The triangle marks Sig 8, whose loadings define the rule shown in (c)–(e). **b,** Per-signature $\hat{\tau}_0$-stratified PFS arm-HR forest. Each row: within-stratum Cox HR (FFX vs GA) with 95% Wald CI; red HR $< 1$, blue HR $> 1$. Right-side column: within-stratum $n$ and two-sided Wald $P$. **c,** Progressive-subgroup PFS arm-HR forest for the Sig 8 triplet rule (TP53 coding mutation; 17p intact, length-weighted log2 $\geq -0.2$; 20q amplified, log2 $> +0.2$). Subgroups ordered top-to-bottom from full triplet to component subsets to "none of the three". Right-side column: within-subgroup $n$ and two-sided Wald $P$. **d, e,** Within-subgroup PFS Kaplan–Meier for patients matching the full triplet (d) and patients matching none of the three criteria (e), by treatment arm. In each KM panel $n$ is the within-stratum size, HR is the Cox FFX-vs-GA estimate, and $P$ is the two-sided Wald $P$-value.

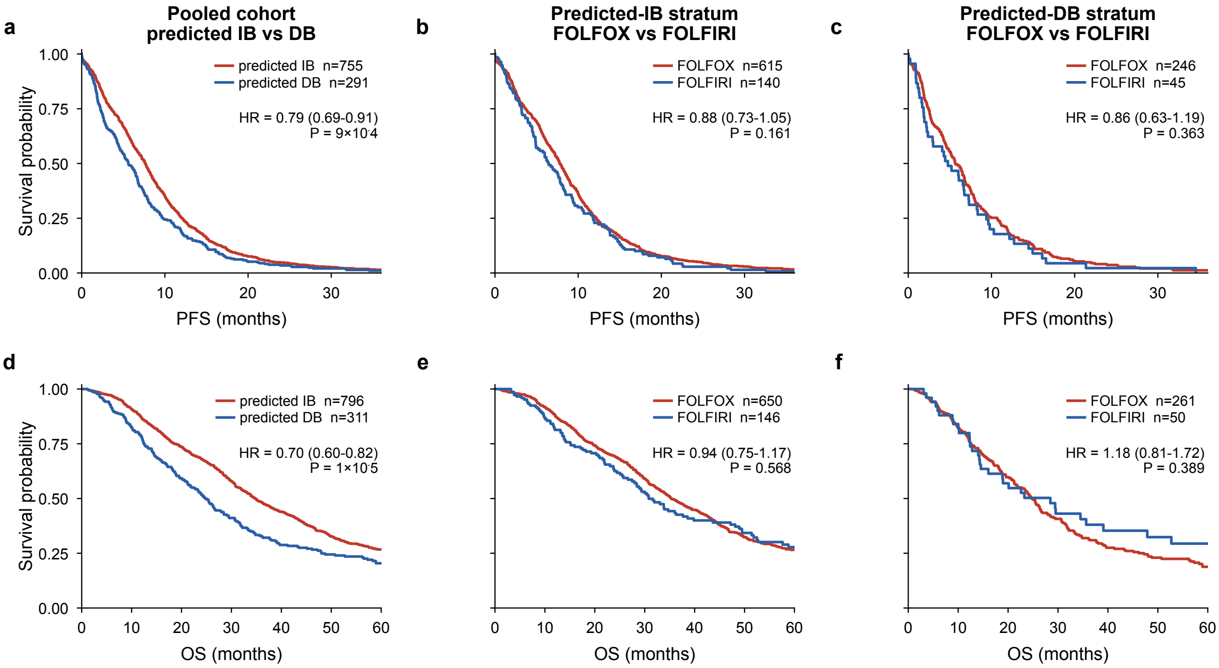


**Supplementary Figure 13. FOLFOXai signature head-to-head on MSK-CHORD.** The FOLFOXai ensemble (five random-forest configurations of 1,000 trees each over a 67-gene panel of somatic mutations and copy-number calls) was re-implemented on the same MSK-CHORD first-line stage IV CRC FOLFOX/FOLFIRI cohort used in Fig. 6 and analysed under the same cut as panels c–e of that figure. Of Abraham's 67 panel genes, 28 (42%) are profiled by MSK-IMPACT and contribute non-zero features; the remaining 39 are absent from MSK-CHORD calls. Abraham's 3% no-call buffer ($0.47 \le p_{\mathrm{IB}} \le 0.53$) excludes 336 of 1,443 patients (23%) from any prediction; the remaining 1,107 patients populate the OS panels below (PFS panels: 1,046 patients with a recorded first-progression date). **a–c,** PFS at 36-month administrative censor. **d–f,** OS at 60-month administrative censor. **a, d,** Pooled cohort split by FOLFOXai predicted class: predicted IB (red) versus predicted DB (blue). **b, e,** Within the predicted-IB stratum, FOLFOX-arm (red) versus FOLFIRI-arm (blue) survival. **c, f,** Within the predicted-DB stratum, arms as in (b). In each panel, $n$ denotes the within-stratum sample size, HR is the Cox proportional-hazards estimate of the first-listed group versus the second-listed group with 95% Wald CI, and $P$ is the associated Wald $P$-value.
